# Mechanisms driving endosperm-based hybrid incompatibilities: insights from hybrid monkeyflowers

**DOI:** 10.1101/461939

**Authors:** Taliesin J. Kinser, Ronald D. Smith, Amelia H. Lawrence, Arielle M. Cooley, Mario Vallejo-Marin, G.D. Conradi-Smith, Joshua R. Puzey

## Abstract

Angiosperm endosperm requires genomic and epigenomic interactions between maternal and paternal genomes for proper seed development. Genomic imprinting, an epigenetic phenomenon where the expression of certain genes is predominantly contributed by one parent, is an essential part of this process and unique to endosperm. Perturbation of imprinting can be fatal to developing seeds, and can be caused by interspecific or interploidy hybridization. However, underlying mechanisms driving these endosperm-based hybridization barriers are not well understood or described. Here we investigate the consequences of genomic imprinting in a naturally occurring interploidy and interspecies hybrid between the diploid, *Mimulus guttatus,* and the allotetraploid (with two subgenomes), *M. luteus* (Phrymaceae). We find that the two parental species differ in patterns of DNA methylation, gene expression, and imprinting. Hybrid crosses in both directions, which suffer from endosperm abnormalities and decreased germination rates, display altered methylation patterns compared to parent endosperm. Furthermore, imprinting and expression patterns appear perturbed in hybrid endosperm, where we observe global expression dominance of each of the two *M. luteus* subgenomes, which share similar expression patterns, over the *M. guttatus* genome, regardless of crossing direction. We suggest that epigenetic repatterning within the hybrid may drive global shifts in expression patterns and be the result of diverged epigenetic/regulatory landscapes between parental genomes. This may either establish or exacerbate dosage-based epistatic incompatibilities between the specific imprinting patterns that have diverged between parental species, thus driving potentially rapid endosperm-based hybridization barriers.

## INTRODUCTION

The evolution of nutritive endosperm is thought to be closely tied to the evolutionary success of angiosperms (W. E. Friedman 1995; Baroux, Spillane, and Grossniklaus 2002). Uniquely, angiosperm seeds are produced by two separate fertilization events (William E. Friedman, Madrid, and Williams 2008). The first event gives rise to the diploid zygote. The second, between an additional sperm cell and two maternal polar nuclei (which are genetically identical to the egg), gives rise to a triploid endosperm in most species. Nutrients and hormones from the maternal plant are transported to the endosperm within the seed, where they nourish and stimulate embryo development. Unlike gymnosperms, in angiosperms, since the endosperm typically cannot develop until fertilized, resources are not wasted on unfertilized eggs (Baroux, Spillane, and Grossniklaus 2002).

After fertilization of the endosperm, genomic interactions between the maternal and paternal genomes control endosperm development and are characterized by genomic imprinting. Genomic imprinting is an epigenetic phenomenon where the alleles of certain genes are regulated differentially depending on their parent of origin, resulting in parent-specific patterns of gene expression (Matzke 1993). RNA and protein production from an imprinted gene is largely restricted to either the maternal allele (termed a maternally expressed gene - MEG) or the paternal allele (paternally expressed gene - PEG) (Kinoshita 2007; Matzke 1993). Genomic imprinting has been observed in a wide range of angiosperm species (Waters et al. 2011; Kinoshita et al. 2004; Florez-Rueda et al. 2016), though our understanding of mechanisms controlling imprinting in plants comes largely from maize and Arabidopsis. Studies in these model systems have revealed that many imprinted genes are associated with differentially methylated regions (DMRs) (Klosinska, Picard, and Gehring 2016; Mei Zhang et al. 2011; Waters et al. 2011; Guo et al. 2003; Hsieh et al. 2009). Prior to fertilization, the maternal alleles of the endosperm are hypomethylated, which may result in differing epigenetic landscapes between the maternal and paternal alleles, resulting in allele specific expression (i.e. imprinting) (Gehring et al. 2006; Ibarra et al. 2012). In Arabidopsis, imprinting status is closely tied to the hypomethylation of transposable elements (TEs) and their resulting DMRs (Gehring, Bubb, and Henikoff 2009). While the function or relevance of many MEGs and PEGs is unknown, several appear to have specific roles. Many MEGs and PEGs in maize and Arabidopsis are associated with epigenetic regulators (Gehring and Satyaki 2016). Certain PEGs are responsible for auxin production, and others are associated with regulating cytokinesis (G. Wang and Köhler 2017), while some regulatory MEGs maintain imprinting states throughout endosperm development (Gehring and Satyaki 2016). Overall, MEGs are generally predicted to restrict endosperm proliferation, and PEGs are predicted to promote it (Gehring and Satyaki 2016). Such regulation in maize and Arabidopsis is essential for proper development of the endosperm and thus the seed.

Traditionally, the establishment and function of parent-specific gene expression in the endosperm is most commonly attributed to opposing strategies for parental and offspring success. Under the parental(kin)-conflict model, conflict arises between the mother and offspring, and between siblings. Selection favors strategies where mothers distribute resources among their offspring to maximize the number that are viable (Haig and Westoby 1989; Haig 2013). Offspring, however, are less related to their siblings in mating systems where they are sired by different fathers, and therefore are selected to acquire more resources than their siblings. It is argued that the paternal filial allele (‘patrigene’) should ‘selfishly’ pull resources into the fertilized seed and the maternal filial allele (‘matrigene’), which is shared among siblings, should repress such demands, allowing resources to be distributed to all related seeds (Haig and Westoby 1989; Haig 2013). Such conflict is proposed to have driven the peculiar two maternal to one paternal genome ratio (2m:1p) in angiosperm endosperm (Haig and Westoby 1989; Haig 2013; Stewart-Cox, Britton, and Mogie 2004), with the idea that it may allow for greater maternal control of resource distribution among seeds (Westoby and Rice 1982).

Departures from the balance of the 2m:1p parental genome dosage results in abnormal endosperm development likely due to misregulation of genomic imprinting and can act as a hybridization barrier (Haig and Westoby 1991a; Leblanc, Pointe, and Hernandez 2002; Pennington et al. 2008). Interploidy crosses, where a tetraploid (*4x*) hybridizes with a diploid (*2x*), alters this 2m:1p ratio, and thus the balance between imprinted patrigene and matrigene (ie. PEG and MEG, respectively) activity. Interestingly, there is a non-reciprocal pattern to this imbalance. When the maternal progenitor is the tetraploid (termed maternal excess; *4x x 2x, maternal x paternal*) the endosperm has increased maternal genomic dosage and thus a 4m:1p ratio. In the reciprocal cross, when the paternal progenitor is the tetraploid (paternal excess; *2x x 4x*), the ratio is 2m:2p (1:1). Such crosses may result in “parent-of-origin” effects, where endosperm size is increased under paternal excess and decreased under maternal excess.

In addition, inter-species hybridization can result in similar asymmetric, non-reciprocal phenotypes even when the species’ ploidies are identical. This is again attributed to departures in the balance between MEGs and PEGs, though in this case, the imprinted “settings” of these genes are expected to have diverged between isolated populations or species (Haig and Westoby 1991). Such ideas have led to hypotheses attempting to explain the role of genomic imprinting as an inter-species hybridization barrier. Extending from observations of inter-ploidy hybrids, the endosperm balance number (EBN) hypothesis predicts that the strength of imprinting differs between imprinted genes of diverged species thus producing “effective ploidy” differences. A diploid species may have stronger imprinting states than another, and thus the hybrid endosperm would experience the same imbalance as an inter-ploid. This hypothesis has been tested in artificial crosses (Johnston and Hanneman 1982) as well as in a naturally-occurring system of *Arabidopsis* species, where incompatibilities at multiple imprinted loci provide a hybridization barrier between two diploid species. This barrier was overcome by the natural tetraploidization of the species with the lower “effective ploidy”, which presumably restored the realized 2m:1p ratio (despite the actual 4m:1p or 2m:2p ratios; (Lafon-Placette et al. 2017)). A similar hypothesis, the weak inbreeder/strong outbreeder (WISO) hypothesis, is specific to imbalances caused by differing mating systems and is a direct extension of the parental-conflict theory. WISO predicts that imprinting should be weaker in highly self-fertilizing species since matrigenes and patrigenes are from the same parent and share siring goals. Thus, hybridization with an outcrossing species would lead to the same imbalances as above (Brandvain and Haig 2005). This hypothesis is also supported with empirical evidence in *Capsella* species (Rebernig et al. 2015) and several other systems (Brandvain and Haig 2005). That said, the mechanisms driving the imbalances that produce these interploidy and interspecies hybridization barriers are not well understood or described.

Various phenomena can drive hybridization incompatibilities and could extend to those occurring within the endosperm. Several recent studies have shown that multiple loci are involved in these endosperm-based hybridization barriers (Rebernig et al. 2015; Lafon-Placette et al. 2017; Garner et al. 2016; Burkart-Waco et al. 2012), which allows for the possibility of Dobzhansky-Muller (DM) like incompatibilities. Under the DM model, a mutation at a given locus may be harmless within the context of the population from which it arose, but when brought together with mutations at interacting loci from a diverged population, incompatibilities can occur (Presgraves 2010). Such negative epistatic interactions have been extended to epigenetic differences (Lafon-Placette and Köhler 2015), including those involving imprinted loci (Josefsson, Dilkes, and Comai 2006). Furthermore, genomic differences between species, particularly regarding TEs, are known to result in genomic shock in hybrids, where structural and regulatory changes may induce negative epistasis and global reprogramming such as subgenome expression dominance (M-J Yoo, Szadkowski, and Wendel 2013; Lafon-Placette and Köhler 2015). Subgenome expression dominance is a phenomenon where the majority of genes from one ‘subgenome’ (the subgenomes are the two newly united parental genomes) are more highly expressed than their corresponding homologs in the other (Mi-Jeong Yoo et al. 2014). Such imbalances in the dosage of interacting components is predicted to drive hybrid incompatibilities (Josefsson, Dilkes, and Comai 2006; Dilkes and Comai 2004). Delineating mechanisms behind hybridization barriers within the endosperm is difficult and is further complicated by the fact that a significant number of plant hybridization events occurring in nature involve multiple ploidies (and thus multiple ‘subgenomes’) and result in variable patterns of asymmetry (Ramsey and Schemske 1998; M. Vallejo-Marin et al. 2016). Elucidating these mechanisms in such systems can help in understanding the role and importance of genomic imprinting in endosperm-based hybridization barriers. These barriers have not only historically been vital in the study of genomic imprinting (Haig and Westoby 1991), but are increasingly recognized to be widespread and major drivers of plant speciation (Lafon-Placette et al. 2017; Lafon-Placette and Köhler 2016).

Here we use a naturally-occurring *Mimulus* hybrid system that is both interploidy and interspecies as a model to explore mechanisms underlying endosperm-based hybridization barriers. *M. guttatus* is a diploid from North America, and *M. luteus*, from the Andes of South America, is an allotetraploid (i.e. formed from past inter-species hybridization and subsequent chromosome duplication) with two distinct subgenomes (subgenomes ‘A’ and ‘B’ (Edger et al. 2017)). Each species is capable of selfing but preferentially outcrosses (Lila Fishman and Willis 2008; Medel, Botto-Mahan, and Kalin-Arroyo 2003). Both were introduced to the British Isles as ornaments in the early 19th century (Parker 1975; Vallejo-Marín et al. 2015); (M. Vallejo-Marin et al. 2016), where they soon naturalized and hybridized producing sterile triploid hybrids (named *M. x robertsii*) (M. Vallejo-Marin and Lye 2013; MarioVallejo-Marin 2012). Most of these hybrids, whether in nature or reproduced in the lab, are formed from *2x x 4x* hybridization events (*M. guttatus* is seed parent; i.e. paternal excess). The reciprocal cross, *4x x 2x* (*M. luteus* is the seed parent – maternal excess) is usually inviable (M. Vallejo-Marin et al. 2016). Such asymmetry is indicative of parent-of-origin effects and offers a unique opportunity to study imprinting in a natural system. One note of phylogenetic clarification -- *Mimulus* as referred to here is non-monophyletic and thus does not represent a true lineage in nature. *M. guttatus* and *M. luteus* are now circumscribed as *Erythranthe* (*E. guttata*, and *E. lutea*, respectively), a well-supported clade (Barker et al. 2012). However, here we refer to them with their commonly used names for communication purposes and to remain consistent with recent work in this hybridization system.

While maize and Arabidopsis have been invaluable models for exploring the mechanisms underlying imprinting, *Mimulus* provides new opportunities for testing these mechanisms in naturally occurring systems. Hybridizations between *Mimulus* species (most now named *Erythranthe* or *Diplacus*) are abundant in nature and have been used to model fundamental questions regarding a diversity of post-zygotic barriers (L. Fishman and Willis 2001; Lila Fishman and Willis 2006; Oneal, Willis, and Franks 2016). There are a variety of life history traits, mating systems, ongoing speciation events, and reticulate patterns (Twyford and Friedman 2015; Grossenbacher and Whittall 2011; Ferris et al. 2017; Brandvain et al. 2014) with which to test hypotheses on the evolution and mechanisms of genomic imprinting. Another unique aspect of this system, distinguishing it from others where imprinting and parent-of-origin effects in endosperm have been characterized (Leblanc, Pointe, and Hernandez 2002; Hehenberger, Kradolfer, and Köhler 2012; Ishikawa et al. 2011; Xu et al. 2014; Rebernig et al. 2015; Florez-Rueda et al. 2016; Meishan Zhang et al. 2016; Scott et al. 1998), is its form of endosperm development. Most angiosperms, including maize and Arabidopsis, undergo nuclear endosperm development, where several rounds of nuclear proliferation precede cytokinesis, whereas *Mimulus* (and other groups within Lamiales) undergo cellular development, where the endosperm proliferates through typical cellular division. While nuclear endosperm development is more common and has evolved repeatedly throughout angiosperms, cellular endosperm development is most likely the ancestral state (Geeta 2003). Importantly, both *M. guttatus* and *M. luteus* have assembled and annotated genomes allowing for detailed genomic analyses (Hellsten et al. 2013; Edger et al. 2017).

In this study we examine patterns of filial tissue development and genomic imprinting in these two *Mimulus* species and their hybrids to investigate mechanisms driving hybridization incompatibilities. First, we use histology to compare seed development patterns and describe the endosperm barrier of reciprocal hybrids in detail. Second, using RNA-seq from developing embryo and endosperm tissues, we define the patterns of imprinting in each species. Third, we determine shifts in imprinting and overall expression patterns within the reciprocal hybrids, where we separate and test relationships between the hybrids’ three subgenomes: the *M. guttatus* genome and the two *M. luteus* ‘A’ and ‘B’ subgenomes. Given that embryo and endosperm are the tissues where genomes contributed by different species first interact, we can more generally investigate subgenome interactions at the earliest stages. Finally, we utilize whole genome bisulfite sequencing of endosperm to characterize differences and changes in DNA methylation of both gene bodies and TEs between the two species and their hybrids. In conclusion, we describe endosperm-based hybrid incompatibilities in the context of subgenome expression dominance and Dobzhansky-Muller like incompatibilities between species-specific imprinted genes.

## RESULTS

### Reciprocal hybrid crosses suffer from developmental abnormalities

To test for abnormalities in seed development of the hybrids, four reciprocal crosses were performed using plants from the CS line for *M. luteus* and plants from the CG line for *M. guttatus* (M. Vallejo-Marin et al. 2016). Crosses are always denoted as *seed parent x pollen donor*. The two hybrid crosses are *M. guttatus* x *M. luteus* (CG x CS, denoted as *2x x 4x*) and *M. luteus* x *M. guttatus* (CS x CG, denoted as *4x x 2x*).

Our first goal was to characterize endosperm and embryo development in *intra*specific crosses. Developing seeds were collected at 3, 5, 8, and 11 days after pollination (DAP), and mature seeds were collected at 15-18 DAP (when fruits dehisced) from *M. luteus* and *M. guttatus* plants. Scanning electron microscopy and histology were used to visualize external and internal seed anatomy, respectively (Fig. 1 and 2). For histology, developing seeds were embedded in resin and serial semi-thin sections were produced with an ultramicrotome. After selecting sections from the center of developing seeds for consistency, whole seed, embryo, and endosperm area was measured (8 and 11 DAP only) using ImageJ. For mature seeds, whole seed area and aspect ratio (AR) was measured. AR is the ratio of width to height, where an AR of 1 is a perfect circle.

**Figure 1.**
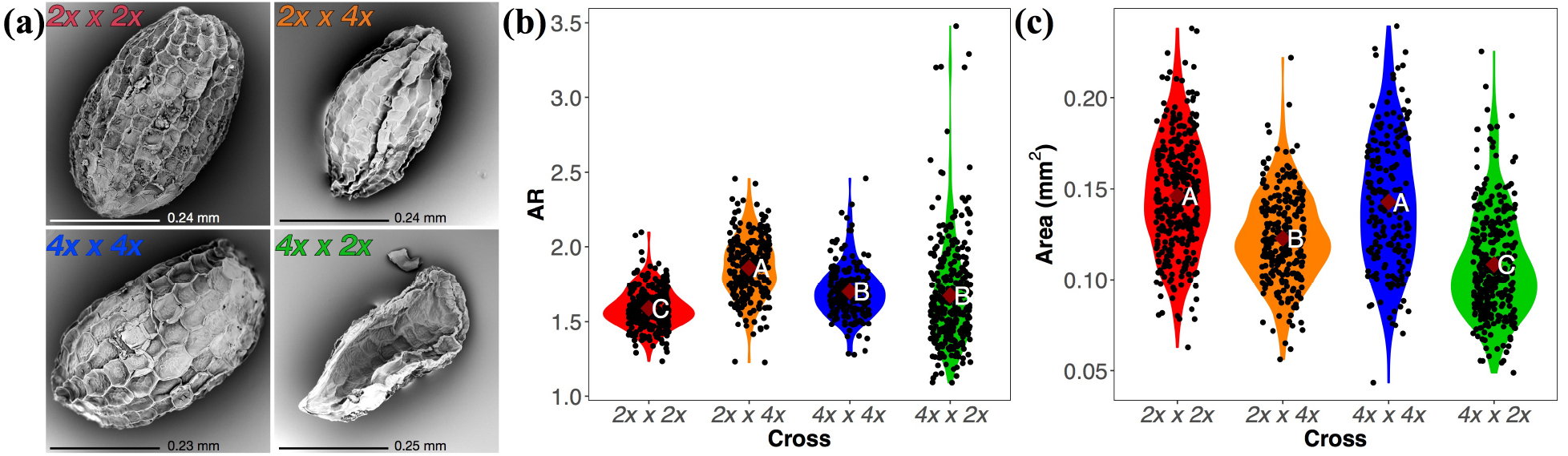
Seed morphology of *M. luteus*, *M. guttatus*, and reciprocal hybrid crosses. Scanning electron micrograph images of a representative seed from each cross (*M. guttatus* – *2x x 2x* [red]; *2x x 4x* [orange]; *M. luteus* – *4x x 4x* [blue]; *4x x 2x* [green]) is displayed in (a). Seed aspect ratio (b) and area (c) in each cross were measured and are represented in violin plots. Dark red diamonds display group means. Letters represent Tukey-Kramer results following an ANOVA. Groups with different letters are statistically different in area or AR.

**Figure 2.**
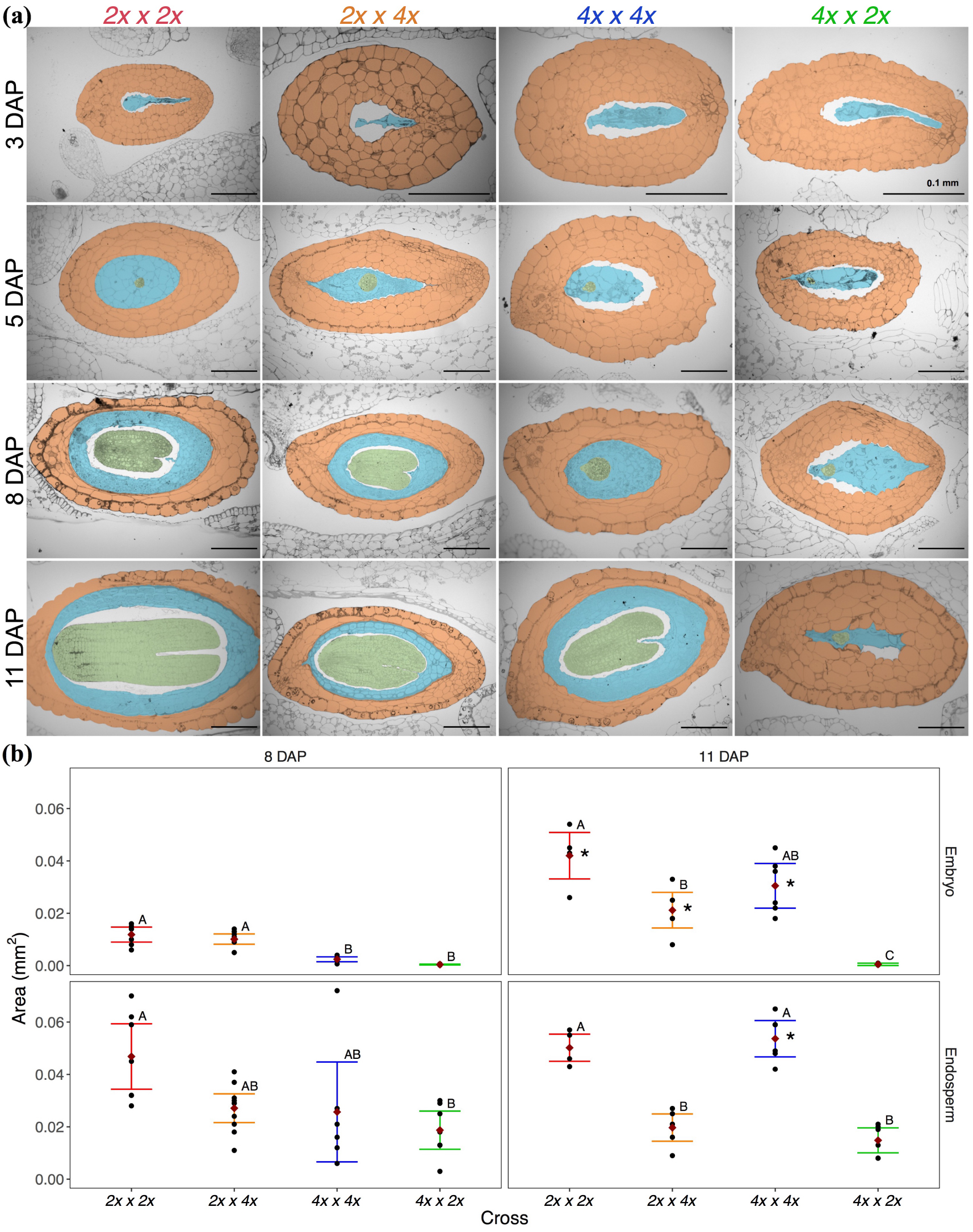
Seed development of parent and reciprocal hybrid seeds. (a) Histological sections were made of each cross through a developmental progression of 3, 5, 8, and 11 days after pollination (DAP). Crosses (*M. guttatus* – *2x x 2x* [red]; *2x x 4x* [orange]; *M. luteus* – *4x x 4x* [blue]; *4x x 2x* [green]) are displayed in columns and DAP is displayed in rows. Within the images, the seed coat is orange, embryo is green, and endosperm is blue. (b) Area of embryo (top) and endosperm (bottom) for all crosses at 8 DAP (left) and 11 DAP (right) was measured. Dark red diamonds represent group means, and error bars represent 95% confidence intervals. Letters represent Tukey-Kramer results following an ANOVA. Groups with different letters are statistically different in area. Asterisks at 11 DAP represent a significant change in tissue size (student’s t-test) from 8 to 11 DAP for the given cross.

At 5 DAP, *M. guttatus* and *M. luteus* embryos were in the globular stage (Fig. 2a). However, by 8 DAP, *M. guttatus* embryos were at a developmental stage between the heart and torpedo stage, while *M. luteus* embryos were smaller and had not yet entered the heart stage (Fig. 2; Supp. Table 3A; Supp. Table 4A). At 11 DAP, both species’ embryos had increased in size (Supp. Table 5) and were in the torpedo stage. While not significantly larger, *M. guttatus* embryos appeared slightly further developed than *M. luteus*’ (Fig. 2; Supp. Table 3B; Supp. Table 4B). Similarly, the endosperm of *M. luteus* developed more slowly than *M. guttatus’* (Fig. 2; Supp. Table 3C; Supp. Table 4C), but both species’ endosperm was large and similar in size by 11 DAP, with *M. luteus*’ showing significant growth from 8 DAP (Fig. 2; Supp. Table 3D; Supp. Table 4D; Supp. Table 5). By maturity, the majority of *M. guttatus* and *M. luteus* seeds were relatively large, round, and plump (Fig. 1a) with high germination rates (Supp. Table 6), and both species’ seeds were similar in area (Fig. 1c; Supp. Table 1B; Supp. Table 2B), though *M. guttatus* seeds were somewhat rounder (Fig. 1b; Supp. Table 1A; Supp. Table 2A).

Next, after describing endosperm and embryo development in intraspecific crosses, we sought to understand developmental differences in the interspecies hybrids. The same methods as above were used for hybrid seeds. The embryos of *2x x 4x* (CG x CS) seeds were in the globular stage at 5 DAP (Fig. 2a) and maintained the developmental progression of their maternal progenitor, *M. guttatus*, through 8 DAP, growing at a faster pace than *M. luteus* embryos (Fig. 2; Supp. Table 3A; Supp. Table 4A). That said, morphological differences from parental endosperm can be observed qualitatively as early as 3 DAP (Fig. 2a). While not significantly different in size at 8 DAP (Fig. 2; Supp. Table 3C; Supp. Table 4C), *2x x 4x* endosperm was smaller than both *M. guttatus’* and *M. luteus’* by 11 DAP (Fig. 2; Supp. Table 3D; Supp. Table 4D) and was not larger than it was at 8 DAP (Supp. Table 5). Similarly, the *2x x 4x* embryo was smaller than *M. guttatus’* at 11 DAP (Fig. 2; Supp. Table 3B; Supp. Table 4B), though it did significantly increase in size from 8 DAP (Supp. Table 5). At maturity, the *2x x 4x* seeds were shriveled (Fig. 1a) and smaller than both *M. guttatus* and *M. luteus* seeds (Fig. 1c; Supp. Table 1B; Supp. Table 2B), and were the narrowest seeds of all the crosses (Fig. 1b; Supp. Table 1A; Supp. Table 2A). The *4x x 2x* (CS x CG) embryo showed no growth from 8 to 11 DAP and was the only cross type that did not reach the torpedo stage of development (Fig. 2; Supp. Table 3A,B; Supp. Table 4A,B; Supp. Table 5). The endosperm also showed little growth and development and was significantly smaller than *M. guttatus* and *M. luteus* endosperm by 11 DAP (Fig. 2; Supp. Table 3C,D; Supp. Table 4C,D; Supp. Table 5). Mature seeds were flattened (Fig. 1a) and were the smallest seeds of all crosses (Fig. 1c; Supp. Table 1B; Supp. Table 2B).

### Endosperm abnormalities are linked to failed or delayed germination of seeds

To test for a link between the morphological abnormalities discussed above and the viability (germination) of the seed, we aligned mature seeds (collected 15-18 DAP) from the four crosses (ie. CG x CG [*M. guttatus*], CS x CS [*M. luteus*], CG x CS [*2x x 4x*], and CS x CG [*4x x 2x*]) along gridded filter paper within petri dishes, photographed them, measured their area in ImageJ, and then recorded their germination date. This allowed us to track the germination success and timing of a given seed and correlate these to its size (area). Since these images were of the whole seed and we could therefore not measure the sizes of internal tissues, we used the histological data above (Fig. 2) to test for relationships between whole seed area and endosperm area. We found that endosperm area has a close relationship with whole seed area (N = 53, R^2^=0.69; Fig. 3a). We therefore consider whole seed area as a good predictor of endosperm area.

**Figure 3.**
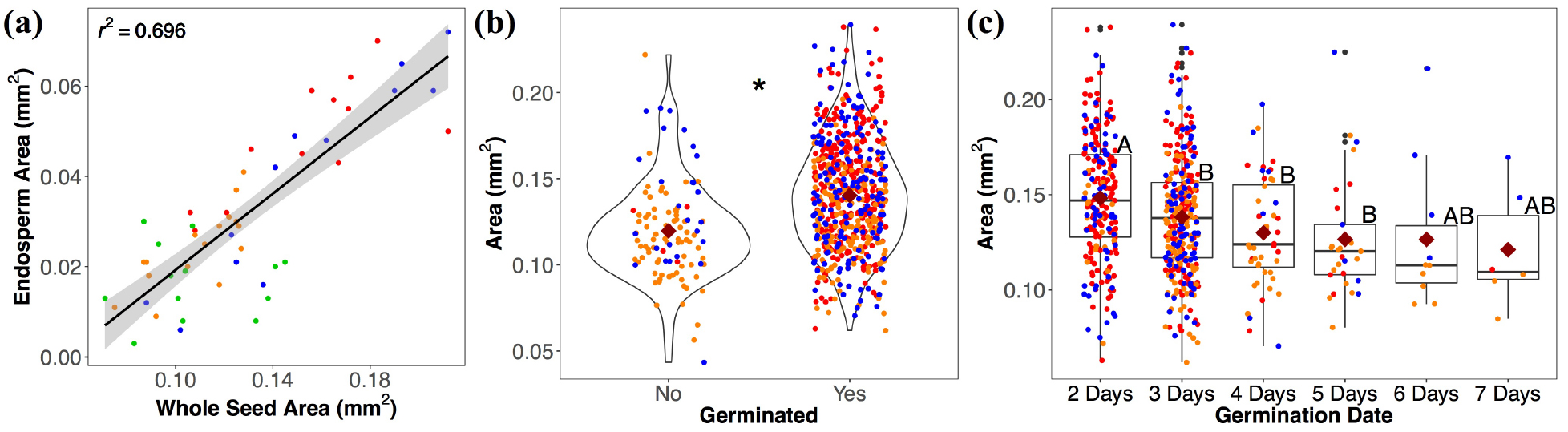
Endosperm relationship to seed area and effect of seed area on germination. (a) Displays a regression between endosperm area and the whole seed area. Colors represent crosses (*M. guttatus* – *2x x 2x* [red]; *2x x 4x* [orange]; *M. luteus* – *4x x 4x* [blue]; *4x x 2x* [green]). Area of seeds that failed to germinate is compared to successfully germinated seeds among all crosses (student’s t-test - significance is represented by the asterisk) in (b). Of seeds that did germinate, their area at the time of germination (2 - 7 days after planting) was compared in (c) with box plots. *4x x 2x* seeds are not included because there were no germinates, and they were the smallest of the four crosses, so as not to bias the comparison. Dark red diamonds in (b) and (c) represent group means. Letters in (c) correspond to Tukey-Kramer tests following an ANOVA. Shared letters indicate that no significant differences were found

The smallest seed type, *4x x 2x*, completely failed to germinate. Across the remaining three seed types, seeds that germinated were significantly larger than seeds that did not (Fig. 3b; Supp. Table 7A, 7B). When considering individual crosses, only *M. luteus* showed no significant difference in size between germinated and ungerminated seeds (Supp. Table 7C-E). Next, we compared the size of seeds that did germinate (which excludes *4x x 2x*) to the timing of their germination. The earliest germinating seeds (2 days after planting) tended to be larger than those that germinated at later dates (Fig. 3c; Supp. Table 8A; Supp. Table 9A). Separate ANOVAs on individual crosses revealed a significant difference among germination dates both for *M. guttatus* and for *2x x 4x*, but subsequent pairwise Tukey’s HSD tests between dates were not significant (Supp. Table 8B-D; Supp. Table 9B-D).

### *Gene expression in the endosperm of* M. luteus *and of* M. guttatus *reveals global paternal bias*

To determine patterns of imprinting and parental bias in *M. luteus*, *M. guttatus*, and the two hybrid crosses (*2x x 4x* and *4x x 2x*), RNA was extracted and sequenced from embryo and endosperm tissue of seeds at the torpedo stage (or equivalent date for *4x x 2x*) in biological triplicate. Imprinted genes were identified by using a reciprocal crossing design to account for line-specific bias. To perform this reciprocal crossing design, two different inbred lines were used each for *M. luteus* (Mll and CS) and *M. guttatus* (CG and LCA). For *M. guttatus*, the two reciprocal crosses are CG x LCA and LCA x CG, and for *M. luteus* the two crosses are Mll x CS and CS x Mll. The reciprocal hybrid crosses are CG x Mll (*2x x 4x*) and Mll x CG (*4x x 2x*).

Whole genome resequencing has previously been conducted on Mll, CS, and CG (M. Vallejo-Marin et al. 2016; Edger et al. 2017), and deep transcriptome resequencing of LCA was conducted in this study. Using these data, line-specific SNPs within genes were identified, allowing RNA-Seq reads generated from endosperm and embryo tissues to be uniquely mapped to specific alleles for measurement of allele expression bias (AEB). AEB is a dimensionless quantity, defined as the mean log_2_[RPKM_paternal_/RPKM_maternal_] across replicates (thus positive AEB denotes higher expression on the paternal allele). RPKM is a normalized metric of gene expression defined as the number of reads per kilobase of coding sequence per million mapped reads (Mortazavi et al. 2008). AEB measurements were standardized so that any amount of expression bias (i.e. AEB does not equal 0) represents a transcriptional departure from the expected genome ratio. Specifically, differences in genomic dosage between the two alleles, due to the 2m:1p parental genome ratio in the endosperm or the tetraploidy of *M. luteus*, was accounted for by adjusting the gene length accordingly in AEB measurements (e.g., for a 2m:1p ratio, the gene length for the maternal allele is multiplied by 2 to represent the doubling of genetic material when calculating RPKM).

To compare the relationships of AEB in the parent species and hybrids, we produced scatterplots of AEB values for genes in one reciprocal cross against their respective values in the other (eg. CG x LCA vs LCA x CG for *M. guttatus*). This plot produced four quadrants representing genes with either consistent line-specific bias (eg. bias towards CG or towards LCA), or consistent maternal or paternal bias (Fig. 4a). Imprinted genes, where allelic expression is consistently biased to either the maternal or paternal allele regardless of crossing direction, should fall into one of the latter two quadrants. To determine whether the consistent parental bias of a gene was truly significant, the likelihood ratio tests developed by Smith, et al. 2017 were used. True imprinted genes are defined here as those that showed a significant shift in bias from one reciprocal cross to the other, thus maintaining strong bias towards maternal (MEGs) or paternal (PEGs) expression.

**Figure 4.**
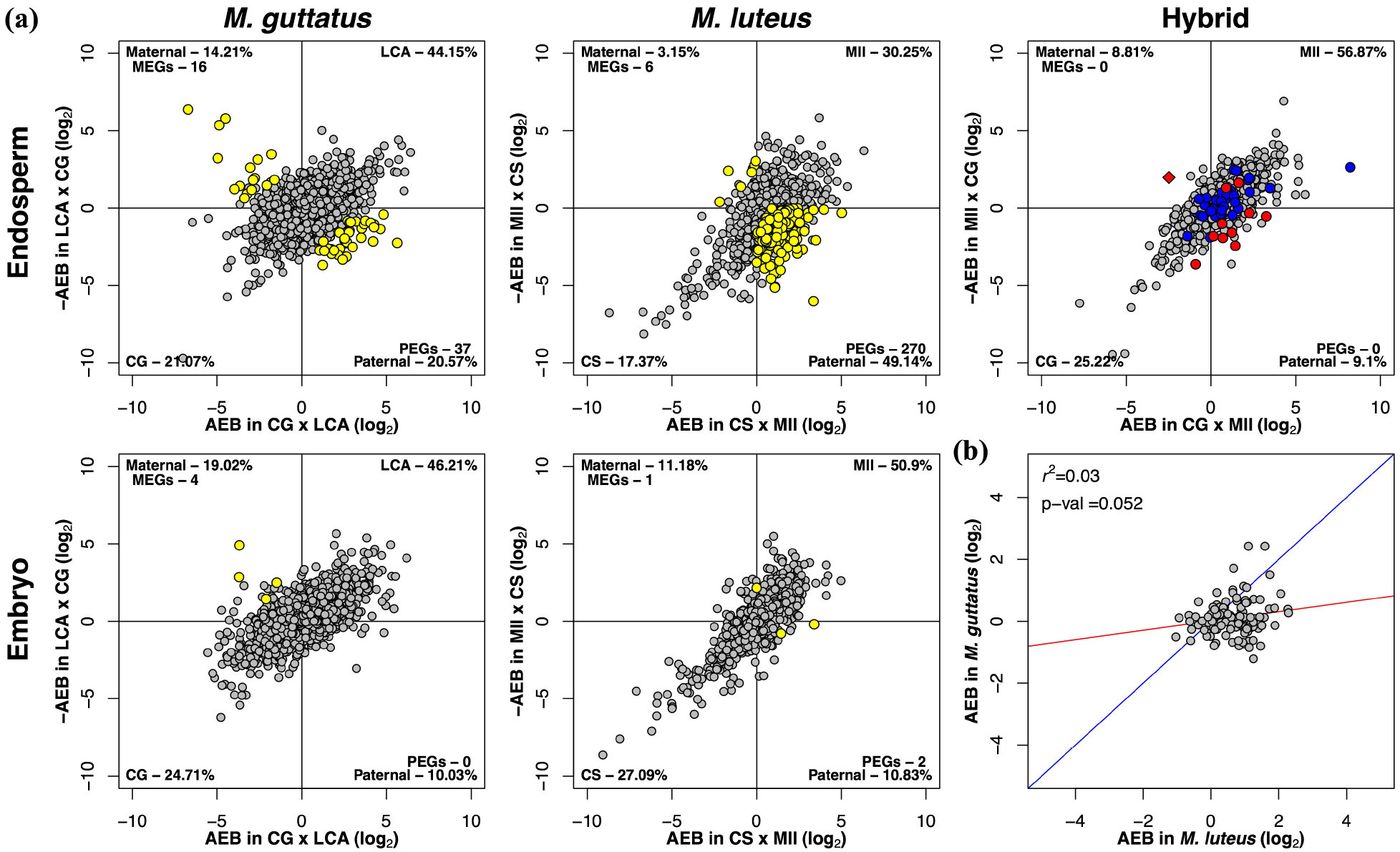
Patterns of imprinting and parental bias. (a) Displays distributions of AEB in the endosperm and embryo of *M. guttatus* and *M. luteus*, and the endosperm of the hybrids. Genes in scatterplots are plotted by their AEB value in one cross on the x-axis against the reciprocal cross on the y-axis. Due to the reciprocal crossing design, bias is divided into four quadrants. Points that fall into the bottom left or top right quadrants represent line or species-specific bias, points in the bottom right have paternal bias, and points in the top left have maternal bias. The percentage of genes in each of these quadrants is given in the corner of that quadrant. The number of PEGs and MEGs is also listed in its respective quadrant. Panels in (a) are sorted by *M. guttatus*, *M. luteus*, and hybrid crosses in columns and endosperm and embryo in rows. A comparison between the hybrids’ embryo is not included since there was insufficient tissue for Mll x CG (*4x x 2x*) embryo. Yellow points signify imprinted genes as determined by a likelihood ratio test (LRT). *M. luteus* and *M. guttatus* PEGs identified within the hybrid endosperm are highlighted in blue and red, respectively. The one MEG from *M. guttatus* is represented by a red diamond. Note these are not significantly imprinted in hybrid endosperm according to LRTs. (b) Compares AEB values between genes in *M. guttatus* and genes in *M. luteus* that are homologs. AEB values of genes are averaged between reciprocal crosses of *M. luteus* and are plotted against AEB values of their homologs in *M. guttatus*, which are also averaged between reciprocal crosses. A linear regression was performed between the homologs of the two species, with the r^2^ value and p-value provided in the top left corner. A hypothetical 1:1 relationship is shown in blue with the actual linear model shown in red.

In the endosperm of intraspecific crosses, 37 putative PEGs and 16 putative MEGs were identified in *M. guttatus*, and 270 PEGs and 6 MEGs were identified in *M. luteus* (Fig. 4a). Overall, 49.14% of genes with any amount of bias fell into the paternal bias quadrant of the AEB scatterplot in *M. luteus* (including those not determined as truly imprinted according to likelihood ratio tests), while only 20.57% of genes did in *M. guttatus* (Fig. 4a; Supp. Table 10). A Fisher test revealed a significantly different PEG:MEG ratio between species (p-value < 0.001), with a greater prevalence of PEGs in *M. luteus*. In contrast to the endosperm, the embryo did not have the same global bias towards paternal gene expression. Only 4 MEGs and no PEGs were identified in *M. guttatus*, while 2 PEGs and 1 MEG were identified in *M. luteus* (Fig. 4a; Supp. Table 10). Overall, we find some MEGs, but many PEGs and global paternal bias in the endosperm of each species, particularly *M. luteus*, and no clear, strong parental bias in either species’ embryo.

### *Gene expression in hybrid filial tissues is dominated by the* M. luteus *genome*

In the hybrid endosperm, we identified no imprinted genes or global parental bias. However, there is a strong overall bias towards *M. luteus*; 56.87% of genes (with any amount of bias) fall into this quadrant (Fig. 4a; Supp. Table 10). Due to stunted *4x x 2x* embryos, we were not able to obtain sufficient material for RNA-seq and could therefore not perform a reciprocal crossing design to distinguish between species and parental bias in the embryos. That said, in the *2x x 4x* embryo, there is also bias towards the paternal progenitor, *M. luteus* (Supp. Table 11). Since angiosperm embryos typically lack imprinting (Gehring and Satyaki 2016), as observed in either parent species’ embryo, this bias likely indicates that the *M. luteus* genome is expressionally dominant in hybrid tissues (for AEB metrics of all parental and hybrid crosses performed in this study, refer to Supp. Table 11). In the following sections, we focus on the endosperm.

### *Patterns of imprinting in the endosperm differ between* M. luteus *and* M. guttatus

To begin to understand how patterns of imprinting may interact between the *M. guttatus* and *M. luteus* genomes, we sought to explore how these patterns are shared between them. We first investigated whether parental expression bias in one species predicts bias in the other by comparing AEB in *M. guttatus* versus *M. luteus*. No relationship in the specific pattern of parental expression bias between homologs of these two species was found (Fig. 4b; Supp. Table 10). Next, we next tested for overlap of imprinted genes between *M. luteus* and *M. guttatus* homologs, though we were limited by their total number of identifiable homologs. Of its 276 putative imprinted genes, *M. luteus* had 38 PEGs with clear homologs in *M. guttatus*. In *M. luteus,* clear homologs were identified for one of *M. guttatus*’ 16 MEGs and five of its 37 PEGs. Note that each of these genes in *M. guttatus* has two corresponding homologs in *M. luteus* since *M. luteus* is a tetraploid. One PEG overlapped between the two species (Mgu_10535 for *M. guttatus*, and Mlu_21243 & Mlu_31004 for *M. luteus* - AT1G51060.1 in Additional Supplement). Finally, we tested for similarities in mean AEB between one species’ set of imprinted genes and the other species’ corresponding homologs. Due to sample size, we only consider PEGs. We measured the overall mean AEB for each species as reference for the mean AEB values of PEGs. These values were averaged between reciprocal crosses to account for line bias and calculated just from this set of *M. luteus* and *M. guttatus* homologs. As expected, in the *M. guttatus* endosperm as well as the *M. luteus* endosperm, even though each have global paternal bias, this bias is stronger for their respective PEGs, especially those of *M. guttatus* (Table 1, Supp. Table 19). In contrast, *M. guttatus* homologs to *M. luteus* PEGs do not share this strong paternal bias within the *M. guttatus* endosperm (we did not have sufficient data for the reverse comparison). Overall, there appears to be substantial differences and limited conservation in imprinting patterns between *M. luteus* and *M. guttatus*.

**Table 1.** Comparison of allele expression bias of PEG homologs between crosses. This table compares AEB values for paternally expressed genes (PEGS) between crosses (*M. guttatus* - *2x x 2x*; *M. luteus* - *4x x 4x*; *2x x 4x*; *4x x 2x*). For an example as to how this table should be read, the mean expression bias of *M. guttatus* homologs of *M. luteus* PEGs is 0.21, indicating little overall expression bias. Genes from one parent species without a homolog identified in the other are not included here. AEB values were averaged between reciprocal crosses (eg. CG x LCA and LCA x CG) for each gene in *M. luteus* (*4x x 4x*) and *M. guttatus* (*2x x 2x*). The overall mean AEB for each cross is listed in the second column. In the third column, the mean AEB of *M. luteus’* PEGs is listed in bold. The mean AEB of the corresponding homologs in *M. guttatus* is reported in the first row. The mean AEB of *M. luteus’* PEGs within *2x x 4x* and within *4x x 2x* is also reported in this column. The first column is similar to the third, but it shows the mean AEB of *M. guttatus’* PEGs in bold and also reports the mean AEB of *M. guttatus’* PEGs within *2x x 4x* and within *4x x 2x*. Note there was not enough data (NED) from the corresponding homologs of *M. guttatus’* PEGs in *M. luteus*. This table shows that expression bias of PEGs is greater than that of the average gene for each parent species and that such expression bias of these PEGs or their homologs is not necessarily maintained or is maintained to a lesser extent in other crosses.

| Cross | Mean AEB of *M. gut* PEGs | Mean AEB Cross | Mean AEB of *M. lut* PEGs |
| --- | --- | --- | --- |
| $2x \times 2x$ (*M. guttatus*) | **2.46** | 0.13 | 0.21 |
| $4x \times 4x$ (*M. luteus*) | NED | 0.64 | **1.30** |
| $2x \times 4x$ (Hybrid) | 1.13 | 0.64 | 0.68 |
| *4x x 2x (Hybrid)* | 1.74 | -0.31 | -0.37 |

### *Within the hybrid endosperm, expression patterns of* M. luteus’ *and* M. guttatus’ *PEGs tend to be consistent with the observed* M. luteus *genome expression dominance*

Next, we tested whether the biased expression of PEGs identified from both *M. luteus* and *M. guttatus* was observed within the hybrids. Within the *2x x 4x* endosperm, on average, expression bias was in the direction of the paternal progenitor, *M. luteus*. Mean AEB values were similar when considering only *M. luteus* PEGs compared to all genes within *2x x 4x* endosperm. However, for PEGs identified in the *M. guttatus* parental cross, their AEB within *2x x 4x* endosperm tended to be more paternally biased than the average gene. That said, *M. guttatus* PEGs were more paternally biased within *M. guttatus* endosperm than they were within *2x x 4x* endosperm (Table 1, Supp. Table 19). Within the *4x x 2x* endosperm, mean AEB was again in the direction of *M. luteus*, the maternal progenitor. *M. luteus* PEGs inherited within *4x x 2x*, reversed to maternal bias. However, *M. guttatus* PEGs maintained paternal bias within the *4x x 2x* endosperm, where *M. guttatus* is the paternal progenitor (Table 1, Supp. Table 19). As evident in the Hybrid Endosperm panel of Fig. 4a, several, but not all, of the *M. guttatus* imprinted genes appear to maintain some level of parental expression bias, whether paternal for *M. guttatus* PEGs or maternal for the one identifiable MEG (though these are not significantly imprinted according to likelihood ratio tests), while many of the *M. luteus* imprinted genes appear to not.

This data suggests that patterns of parental bias are not obviously inherited from either parent species in the hybrids. Rather, while there may be some consistent parental bias from *M. guttatus’* imprinted genes, expression bias within hybrid endosperm appears to remain more consistent with global *M. luteus* expression dominance than with normal patterns of imprinting observed in parent endosperm, especially that of *M. luteus*.

### *In* M. luteus *and* M. guttatus*, maternal expression is down-regulated and paternal expression is up-regulated in endosperm alleles compared to the embryo*

For a more specific understanding of how parental alleles are expressed in intraspecific *M. luteus* and *M. guttatus* crosses, we compared expression (measured as log(RPKM)) across all genes for each allele in endosperm and embryo. Gene expression values were averaged between reciprocal crosses to account for line bias. Next, we used an ANOVA with subsequent pairwise Tukey’s HSD tests to compare overall expression levels among each allele (ie. paternal and maternal) within the embryo and endosperm of both *M. luteus* and *M. guttatus*.

Expression levels did not significantly differ between the maternal and paternal alleles of the *M. luteus* embryo. The paternal allele of the endosperm had significantly higher expression levels than either of the embryo alleles, while the maternal allele of the endosperm was significantly lower than either embryo allele (Supp. Table 18C; Supp. Table 17C; Supp. Fig. 2C). Unlike in *M. luteus,* expression levels of the maternal and paternal alleles in the *M. guttatus* embryo were significantly different. However, as with *M. luteus*, the paternal allele of the endosperm had significantly greater and the maternal allele had significantly lower expression levels than either of the embryo alleles (Supp. Table 18A; Supp. Table 17A; Supp. Fig. 2A).

Finally, in the embryo of *2x x 4x*, expression of the *M. guttatus* genome (ie. maternal allele) was significantly less than that of the *M. luteus* genome (paternal). For the *M. luteus* genome, gene expression did not significantly differ between the embryo and the endosperm of *2x x 4x*, but the *M. guttatus* genome had even lower expression in the endosperm (Supp. Table 18B; Supp. Table 17B; Supp. Fig. 2B). In *4x x 2x* endosperm, gene expression was also greater for the *M. luteus* genome (ie. maternal) than the *M. guttatus* genome (paternal) (Supp. Table 17D; Supp. Fig. 2D).

Overall, these data provide more evidence that the *M. luteus* genome is expressionally dominant in hybrid seeds and suggest that paternal expression is up-regulated and, to a greater extent, maternal expression is down-regulated in *Mimulus* endosperm.

### *The gene bodies of* M. luteus *and* M. guttatus *endosperm differ in their DNA methylation patterns*

To analyze DNA methylation patterns in *M. luteus*, *M. guttatus*, and their hybrids, DNA was extracted from endosperm tissue of seeds at the torpedo stage (or equivalent date for *4x x 2x*) using the CTAB method (Porebski, Bailey, and Baum 1997) and treated with bisulfite for Methyl-Seq. The following four crosses were used: CG x LCA (for *M. guttatus*), Mll x CS (for *M. luteus*), CG x Mll (*2x x 4x*), and Mll x CG (*4x x 2x*). To identify patterns of gene body methylation, reads from CpG, CHG, and CHH sequence contexts were mapped to coding regions on each allele within each cross, and methylation was called using the methylpy pipeline (Schultz et al. 2015). The fraction of methylated CpG, CHG, and CHH sequences was calculated for each gene. The empirical cumulative distribution function (eCDF) and Kolmogorov-Smirnov (KS) tests were used to compare the fraction of methylation on alleles between and within crosses for each sequence context. Gene counts, median fraction of methylation values, and outcomes of KS tests are reported in Supp. Table 12, Supp. Table 13, and Supp. Table 14, respectively. For visualization, heat maps displaying the fraction of methylation on paternal and maternal alleles are provided in Supplemental Figure 1.

Patterns of gene body methylation were first analyzed for the endosperm of parental crosses. Both the maternal and paternal alleles of *M. luteus* had higher levels of CHG and CHH methylation than *M. guttatus*. That said, for both species, most genes have low CHG and CHH methylation levels (Fig. 5a,b; Supp. Table 12A; Supp. Table 13A; Supp. Table 14B-D). CpG methylation was bimodally distributed for both species with the fraction of methylation on a given gene typically near 0 or near 1 (Fig. 5a,b; Supp. Fig. 1). However, a larger portion of genes (on either allele) in *M. guttatus* were highly methylated than in *M. luteus* (Supp. Table 15A; Supp. Figure 1) and thus overall methylation was greater for *M. guttatus* (Fig. 5a,b; Supp. Table 12A; Supp. Table 13A; Supp. Table 14B-D). CpG methylation was higher on the paternal allele than the maternal in *M. guttatus*, but, while statistically different, median values for CHH and CHG are very similar between parental alleles (Fig. 5a,b; Supp. Table 12A; Supp. Table 13A; Supp. Table 14A). There are no statistical differences between parental alleles for any type of methylation in *M. luteus* (Fig. 5a,b; Supp. Table 12A; Supp. Table 13A; Supp. Table 1 4A).

**Figure 5.**
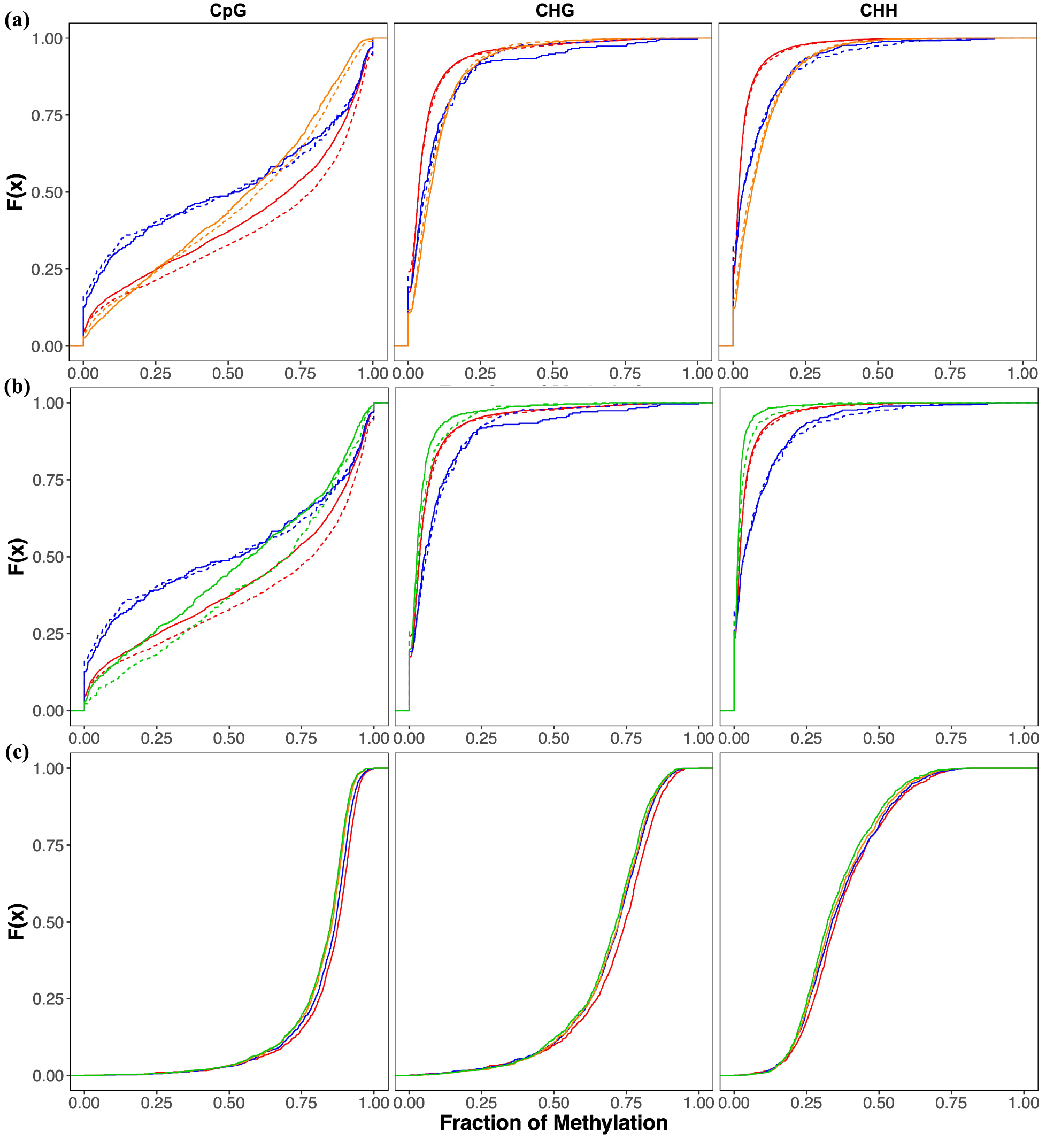
Patterns of methylation in the parents and hybrids. The empirical cumulative distribution function is used to compare levels of methylation (for CpG, CHG, and CHH sequence types) in CG x LCA (for *M. guttatus* – *2x x 2x* [red]), Mll x CS (for *M. luteus* – *4x x 4x* [blue]), CG x Mll (*2x x 4x* [orange]), and Mll x CG (*4x x 2x* [green]). (a) and (b) display gene body methylation on the maternal (solid line) and paternal (dashed) alleles. CG x Mll (a) and Mll x CG (b) are plotted beside *M. guttatus* and *M. luteus*. Methylation on transposable elements is displayed for all crosses in (c). Parental alleles are not differentiated.

### Gene body methylation is repatterned in the hybrid endosperm

Next, patterns of gene body methylation were analyzed for DNA from the endosperm of hybrid individuals. Among all the crosses, *4x x 2x* had the lowest overall fractions of CHG and CHH methylation on both its maternal and paternal alleles (Fig. 5b; Supp. Table 12A; Supp. Table 13A; Supp. Table 14B-D). In contrast, *2x x 4x* had the highest level of CHG and CHH methylation on either allele (Fig. 5a; Supp. Table 12A; Supp. Table 13A; Supp. Table 14B-D). Methylation tended to be higher on paternal than on maternal alleles in *4x x 2x* (Fig. 5b; Supp. Table 12A; Supp. Table 13A; Supp. Table 14A), but higher on maternal than paternal alleles in *2x x 4x* (Fig. 5a; Supp. Table 12A; Supp. Table 13A; Supp. Table 14A). In other words, more CHG and CHH sequences are methylated on the *M. guttatus* allele in the hybrids. That said, the fraction of CHG and CHH gene body methylation was near 0 for most genes (Supp. Table 13A). CpG methylation was less bimodally distributed in *2x x 4x* and *4x x 2x* endosperm than it was in the parental endosperm (Fig. 5a,b; Supp. Fig. 1). Instead, the concentration of genes with a high fraction of methylated CpG sequences was lower in the hybrids than in the parents, as visible in the eCDF tails (note that hybrid eCDF curves are more linear), but an overall larger portion of genes had some CpG sequences methylated in the hybrids than in the parents (Supp. Table 15A). A similar pattern can be observed with CHG and CHH methylation in *2x x 4x*, though it is less notable since this methylation is minimal in the gene bodies (Supp. Fig 1; Supp. Table 15B,C). In agreement with this pattern, median CpG methylation values for maternal and paternal alleles of *2x x 4x* and *4x x 2x* fell between the median values of *M. luteus* and *M. guttatus* (Supp. Table 13A). CpG methylation was higher on the paternal allele than the maternal allele for both *2x x 4x* and *4x x 2x* (Fig5a,b; Supp. Table 12A; Supp. Table 13A; Supp. Table 14A); though this disparity is even greater for *4x x 2x*, where the paternal allele was strongly methylated (Fig5b; Supp. Table 13A). Taken together, this data suggest that methylation patterns depart from those observed in parent species in unique ways for each hybrid and that methylation specificity may be decreased.

### Transposon methylation changes in hybrid endosperm

Using the same Methyl-Seq data as above, reads from CpG, CHG, and CHH sequence contexts were mapped to a curated database of *Mimulus* transposons (TEs) (Edger et al. 2017) and methylation was analyzed using methylpy to identify patterns of TE methylation. Among all the crosses, *M. guttatus* had the highest fraction of TE methylation for all methylation types (Fig5c; Supp. Table 12B; Supp. Table 13B; Supp. Table 14E). Both *2x x 4x* and *4x x 2x* had the least amount of CpG methylation on TEs (Fig5c; Supp. Table 12B; Supp. Table 13B; Supp. Table 14E). The fraction of CHG methylation on TEs did not significantly differ between *M. luteus*, *2x x 4x*, or *4x x 2x*. *M. luteus* had higher levels of CHH methylation than *4x x 2x*, but *2x x 4x* did not significantly differ from either of them (Fig5c; Supp. Table 12B; Supp. Table 13B; Supp. Table 14E). Overall, TE methylation is highest in *M. guttatus* and tends to be lower in the hybrids.

### *Epigenetic patterns tend to be shared between* M. luteus *subgenomes*

Next, we sought to determine patterns of allele expression bias specific to each of the two *M. luteus* subgenomes (A and B subgenomes) within developing *M. luteus* seeds. The homolog to a gene from one *M. luteus* subgenome (eg. A) on the other *M. luteus* subgenome (eg. B) is referred to as a hom*e*olog, and the pair is referred to as a hom*e*olog pair. We first compared overall expression levels (log(RPKM)) among subgenome-specific alleles (subgenomes A and B will be subscripted by ‘m’ or ‘p’ indicating maternal or paternal genome: A_m_, A_p_, B_m_, B_p_) using ANOVAs with subsequent pairwise Tukey’s HSD tests (as described above). We found no significant difference in overall expression among any of the alleles (ie. A_m_ vs. A_p_ vs. B_m_ vs. B_p_) within the embryo (Supp. Table 18D; Supp. Table 17E; Supp. Fig. 2E). In the endosperm, the maternal alleles from separate subgenomes (A_m_ vs. B_m_) shared similar expression levels, and the paternal alleles (A_p_ vs. B_p_) also shared similar expression levels with each other. However, maternal alleles had significantly lower expression levels compared to paternal alleles, regardless of subgenome (A_m_ and B_m_ vs. A_p_ and B_p_)(Supp. Table 18D; Supp. Table 17E; Supp. Fig. 2E).

Next, we compared patterns of parental expression bias (ie. AEB) between subgenomes (ie. *A_m_ x A_p_* and *B_m_ x B_p_*). Within the endosperm, the parental alleles of subgenome A (*A_m_ x A_p_*) show overall paternal expression bias as do the parental alleles of subgenome B (*B_m_ x B_p_*) (Supp. Table 11B). That said, when comparing AEB across individual homeolog pairs (eg. *Gene1-A_m_ x Gene1-A_p_* vs. *Gene1-B_m_ x Gene1-B_p_*, etc.), we found no relationship between the AEB of alleles on one subgenome and that of their homeologs on the other in either the embryo or endosperm (Fig. 6a; Supp. Table 16; Supp. Fig. 3a). Taken together, even though the AEB of a specific gene on subgenome A may differ from its homeolog on subgenome B, the mean AEB across all subgenome A is similar to that of subgenome B.

**Figure 6.**
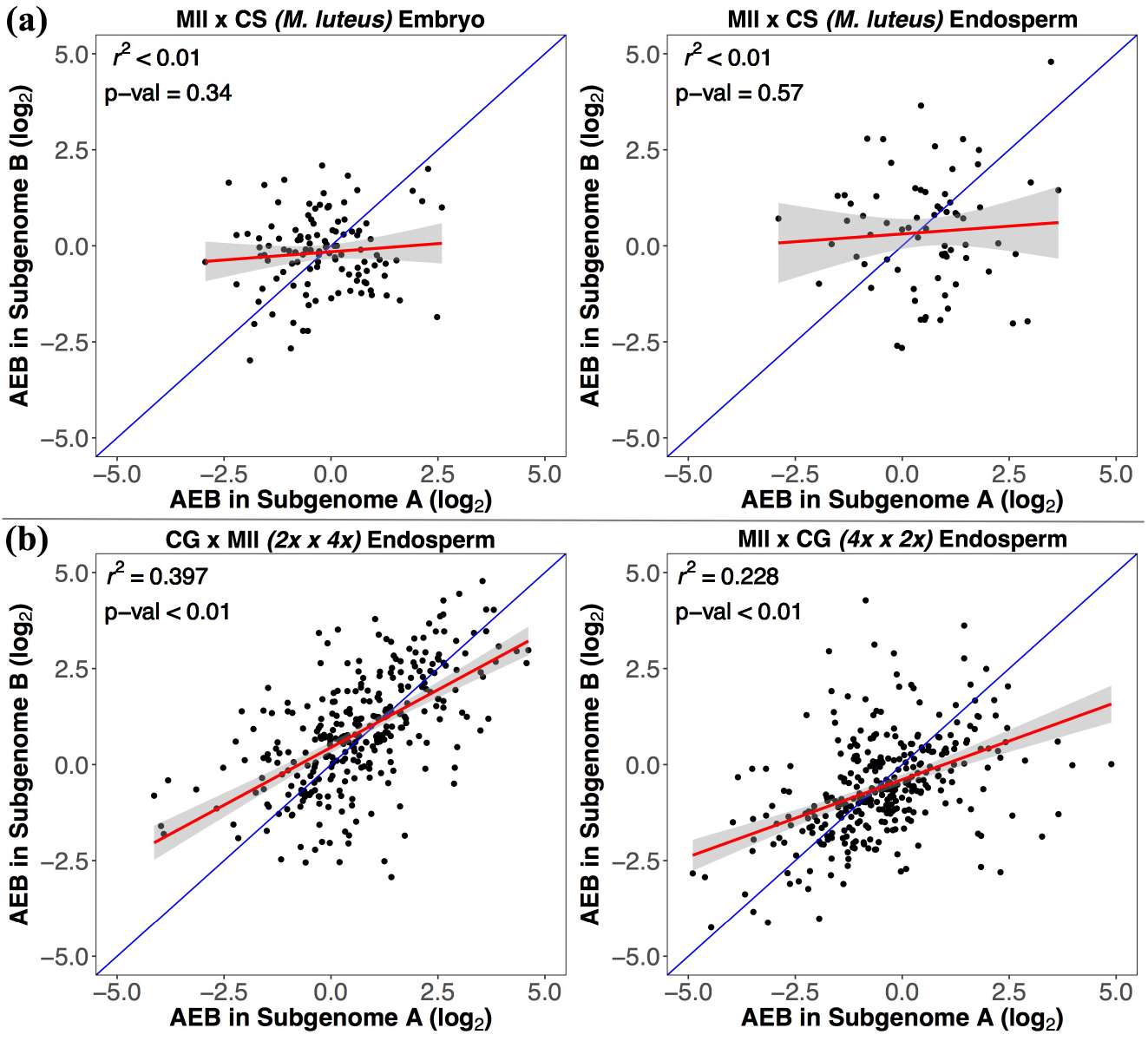
Comparison of AEB values between *M. luteus* subgenomes. AEB values (between maternal and paternal alleles) for genes on *M. luteus’* subgenome A are compared to AEB values of their homeologs on subgenome B. (a) represents subgenomes within seed tissues of *M. luteus* (embryo on the left, endosperm on the right; values for Mll x CS are shown here), and (b) represents subgenomes within the endosperm of hybrid crosses (*2x x 4x* [CG x Mll] on the left, *4x x 2x* [Mll x CG] on the right). R^2^ values and p-values for linear regressions are provided in the top left corner of each plot. A hypothetical 1:1 relationship is shown in blue with the actual linear model in red. The gray shaded area represents the 95% confidence interval of the linear model.

We additionally analyzed expression patterns of the two *M. luteus* subgenomes inherited within hybrid seeds. Global expression levels (log(RPKM)) did not significantly differ between the *M. luteus* subgenomes (paternal) within *2x x 4x* (A_p_ vs. B_p_) in either the embryo or the endosperm (Supp. Table 18E; Supp. Table 17F; Supp. Fig. 2F). Furthermore, there was no significant difference between *M. luteus* subgenomes (maternal) within the *4x x 2x* endosperm (A_m_ vs. B_m_) (Supp. Table 18F; Supp. Table 17G; Supp. Fig. 2G). However, the expression levels of each *M. luteus* subgenome was greater than that of the *M. guttatus* genome in both *2x x 4x* (*M. guttatus*_m_ vs. A_p_ and *M. guttatus*_m_ vs. B_p_) and *4x x 2x* (A_m_ vs. *M. guttatus*_p_ and B_m_ vs. *M. guttatus*_p_) endosperm. Overall expression bias (AEB) towards *M. luteus* was observed for both A (ie. M. guttatus*_m_ x A_p_*) and B (M. guttatus*_m_ x B_p_*) subgenomes in *2x x 4x* embryo and endosperm, as well as in *4x x 2x* endosperm (ie. *A_m_ x* M. guttatus*_p_* and *B_m_ x* M. guttatus_*p*_)(Supp. Table 11B). Furthermore, while *r*^2^ values were somewhat low for *4x x 2x*, there were significant relationships of AEB between homeolog pairs within the embryo and endosperm of *2x x 4x* (eg. *Gene1-*M. guttatus*_m_ x Gene1-A_p_* vs. *Gene1-*M.guttatus*_m_ x Gene1-B_p_*, etc.) and the endosperm of *4x x 2x* (eg. *Gene1-A_m_ x Gene1-*M.guttatus*_p_* vs. *Gene1-B_m_ x Gene1-*M.guttatus_*p*_, etc.). This suggests that expressional biases against the *M. guttatus* allele were consistent between homeolog pairs of the two *M. luteus* subgenomes within the hybrids (Fig. 6b; Supp. Table 16; Supp. Fig. 3b). Thus, expression patterns appear to be more similar between the *M. luteus* subgenomes than with the *M. guttatus* genome within the hybrid.

### Putative functions of imprinted genes

As a first look into the function of imprinting in *Mimulus* endosperm, we implemented a BLAST search of the *M. guttatus* sequence against *Arabidopsis thaliana* on our list of *M. guttatus* imprinted genes. A complete list of all putative imprinted genes (including from *M. luteus*) with full descriptions and *Mimulus* gene IDs is provided in the Supplemental Materials.

Several putative MEGs are associated with or occur in plastids. These include the photosystem II reaction center and genes involved in the response to and redistribution of light energy between photosystems (Xue et al. 2015; Depège, Bellafiore, and Rochaix 2003). Some genes operate in plastids and have broad functions essential to seed development. One such MEG is an intermediary in biosynthesis of abscisic acid (ABA), which establishes seed dormancy and is involved in many other important processes in seeds (Finkelstein, Gampala, and Rock 2002). Another is a 6-phosphogluconate dehydrogenase protein, which are components of the pentose phosphate pathway and important for stress tolerance and seed growth (Spielbauer et al. 2013; Hou et al. 2007). Other MEGs are involved in desiccation and cold tolerance (Kleinwächter et al. 2014; Borovskii et al. 2002) and other stress tolerances (Liu et al. 2015; Roy and Wu 2002), vacuolar storage (Bolte et al. 2011; Jauh, Phillips, and Rogers 1999), metabolism and recycling, cell division and growth, RNA binding, and there is also a ribosomal protein.

Identified PEGs appear to have a variety of functions, particularly related to phytohormone signaling. Along with ABA, other important hormones in seeds include jasmonic acid (JA), which plays a role in herbivory and abiotic stress response, germination, and other developmental processes (Singh et al. 2017; Delker et al. 2006; Yildiz, Muradoglu, and Yilmaz 2008; Creelman and Mullet 1995), and cytokinins, which, among other functions, are involved in seed size and germination (Riefler et al. 2006). Among the identified PEGS, there is a MYC transcription factor and a target of a MYB transcription factor, which both play important developmental roles in response to ABA and JA (Ambawat et al. 2013; Boter et al. 2004; UniProt Consortium 2018) (https://www.uniprot.org/uniprot/Q39204). MYC is additionally involved in light response, also important for seed germination and development (Yan et al. 2014). Another putative PEG is a member of the Protein phosphatase 2C family, which are key in ABA signal transduction (Rodriguez 1998). There is also a member of F-box proteins that is involved with negative regulation of both cytokinin response and phenylpropanoid biosynthesis (Kim et al. 2013; X. Zhang, Gou, and Liu 2013). Finally, we find an ABC transporter, which are ATP-hydrolysing membrane pumps and are involved in many processes, including seed development and transport of auxin and abscisic acid (Kang et al. 2011). Other putative PEGs are involved in bacterial defense and potyviral replication (Duprat et al. 2002; Lee, Jelenska, and Greenberg 2008), protein degradation, vesicle trafficking (UniProt Consortium 2018) (https://www.uniprot.org/uniprot/Q9LK09), and cell wall modification (Table 2). In addition, there is a molecular chaperone protein, an ATPase, a ribosomal protein, and a histone protein (Table 2). This histone protein, histone H2A in *A. thaliana*, is the only shared imprinted gene between *M. guttatus* and *M. luteus* that we could identify. It should be noted, though, that in *A. thaliana*, this gene’s mRNA is cell-to-cell mobile (The Arabidopsis Information Resource, www.arabidopsis.org). This putative histone PEG, conserved between *M. guttatus* and *M. luteus*, may be important for the observed epigenetic patterns in the *Mimulus* endosperm, though it is possible it is carried over from DNA packaging in pollen nuclei. Several other PEGs are expressed in pollen and thus their imprinted status and role in seed development should be verified in future studies. In summary, we find diverse roles in development for both MEGs and PEGs.

## DISCUSSION

Interspecific hybridization brings together two genomes that each have unique evolutionary histories. Studying how these genomes interact at the earliest stage, during embryo and endosperm development, will increase our understanding of not only basic plant processes, such as genomic imprinting and seed development, but also mechanistic barriers to hybridization. Here, using a *Mimulus* hybrid system, which is both interspecies and interploidy, we test for developmental and genomic consequences of hybridization in endosperm and embryos. The comparable endosperm phenotypes and crossing asymmetries observed between interploidy and interspecies hybrids are often linked to similar mechanisms involving aberrant imprinting (Gehring and Satyaki 2016), which are traditionally attributed to a departure from a delicate balance between parental strategies driven by parental conflict (Haig 2013). Interestingly, such hybridizations and their associated parent-of-origin effects are substantially variable, both in crossing asymmetries (M. Vallejo-Marin et al. 2016; Ramsey and Schemske 1998) and underlying changes in expression patterns (Vrana et al. 1998; Josefsson, Dilkes, and Comai 2006; L. Wang et al. 2018; Florez-Rueda et al. 2016; Ishikawa et al. 2011). While theory posed by the endosperm balance number (EBN) and weak inbreeder/strong outbreeder (WISO) hypotheses may help describe general patterns in endosperm-based hybridization barriers, the underlying mechanisms behind many of these barriers are likely more complex than predictable, uniform departures from the genomic balance of a shared set of imprinted loci. This study helps shed light on potentially fundamental mechanisms that may drive these incompatibilities within hybrid endosperm.

In *Mimulus* we found a positive relationship between endosperm size and germination success; and that hybrid seeds have smaller endosperm. The non-reciprocal phenotypes of flat, inviable seeds when *M. luteus* was the seed parent (*4n x 2n*), and shriveled seeds with moderate viability when *M. guttatus* (*2n x 4n*) was the seed parent, are reminiscent of non-reciprocal endosperm abnormalities and seed viability in other interspecies hybrids (Rebernig et al. 2015; Haig and Westoby 1991; Lafon-Placette et al. 2017; Scott et al. 1998). In species with nuclear endosperm development, these abnormalities are usually associated with the timing of cellularization, which is under imprinted control (Ishikawa et al. 2011). While cellular endosperm developmental processes are different, they are also likely under imprinted control since the *Mimulus* hybrid crosses exhibit parent-of-origin effects in their endosperm. Some imprinted genes in *M. guttatus* were involved in cell division and growth and cell wall modification – likely important for cellular endosperm development. The observed parent-of-origin phenotypes - flat and shriveled seeds - may be unique to cellular endosperm species as similar seed phenotypes occur in other *Mimulus* hybrids (Oneal, Willis, and Franks 2016).

We also observed clearly diverged imprinting patterns between *M. guttatus* and *M. luteus*. Rapid evolution and divergence of species-specific imprinting patterns is common and is likely involved in endosperm-based hybridization barriers. Shifts in expression patterns are also common in hybrid endosperm and can directly affect the dosage of imprinted genes and other developmental genes linked to non-reciprocal abnormalities (Klosinska, Picard, and Gehring 2016; Florez-Rueda et al. 2016; Ishikawa et al. 2011; Burkart-Waco et al. 2015; Li and Dickinson 2010). In our system, neither reciprocal hybrid had global imprinting patterns consistent with either parent. Instead we observed consistent subgenome expression dominance of the inherited alleles from *M. luteus* over *M. guttatus*. Furthermore, PEG status, particularly that of *M. luteus* PEGs, was not necessarily maintained in the hybrids. We observed unique DNA methylation patterns of endosperm not only between *M. luteus* and *M. guttatus*, but also within each of the reciprocal hybrids. Such changes in methylation patterns have been shown to alter dosage and drive incompatibilities with typical parent-of-origin phenotypes (Kirkbride et al. 2015; Bushell, Spielman, and Scott 2003).

These results, along with observations made in other systems, lead us to make the following general conclusions of this *Mimulus* hybrid system: (1) We suspect that the endosperm is the primary barrier in seed development due to (a) striking similarities with other systems as discussed above, (b) lack of genomic imprinting in the embryo of either parent species and exaggerated expression dominance in hybrid endosperm compared to hybrid embryo, and (c) past work in this system showing that germinated hybrid seeds produce vigorous, healthy plants (Vallejo-Marín et al. 2016); (2) The specific genes that are imprinted in the endosperm vary between these closely related species; (3) The magnitude of genomic imprinting in the endosperm (overall amount of parental specific expression bias), and thus allele-specific dosage differs between these species; (4) Imprinting patterns are not maintained in the hybrids, and a pattern of subgenome expression dominance emerges; (5) Methylation patterns differ between parents and are altered in developing endosperm of hybrid plants; (6) Hybrid incompatibility may be related to a combination of (a) parental species having different imprinting and general epigenetic patterns and (b) subgenome expression dominance in interspecies hybrids leading to exaggerated differences in gene expression in developing filial tissues. Slight differences in developmental timing of parental species may also affect compatibility (Oneal, Willis, and Franks 2016), though, while it may be under imprinted control, we do not test timing here. Below we expand on these results by placing them in the context of a Dobzhansky-Muller (DM) like incompatibilities and genomic shock.

### Differing epigenetic landscapes between species can drive epigenetic reprogramming in their hybrid, which, coupled with Dobzhansky-Muller like effects on imprinted genes, can act as an endosperm-based hybridization barrier

Here, we observed differences between *M. guttatus* and *M. luteus* in both their sets of imprinted genes and their relative allelic dosages. If different genes are imprinted in parental species, or their imprinting patterns differ in dosage, then when brought together, interactions involving multiple imprinted loci may be mismatched between parental genomes (Yadegari et al. 2000; Josefsson, Dilkes, and Comai 2006). This could result in epigenetic DM-like incompatibilities (Josefsson, Dilkes, and Comai 2006; Lafon-Placette and Köhler 2015; Wolf, Oakey, and Feil 2014); (Garner et al. 2016; Rebernig et al. 2015). Compounding these issues, genomic alterations and epigenetic reprogramming (i.e. genomic shock (McClintock 1984)) often occur during hybridization. Epigenetic reprogramming may cause global shifts in expression patterns, such as subgenome expression dominance (Mi-Jeong Yoo et al. 2014; M-J Yoo, Szadkowski, and Wendel 2013), and could enforce or exacerbate DM-like imprinting incompatibilities (Comai et al. 2003). For example, a common feature in interspecies hybrids is loss, shifting, or gain of imprinting status (Burkart-Waco et al. 2015; Kirkbride et al. 2015; Josefsson, Dilkes, and Comai 2006; Vrana et al. 1998). A combination of these related epigenetic mechanisms could potentially help explain the wide variability of outcomes in hybrid seeds and endosperm (Ishikawa et al. 2011; Garner et al. 2016; Josefsson, Dilkes, and Comai 2006; Rebernig et al. 2015; Florez-Rueda et al. 2016; Burkart-Waco et al. 2015). Below, we outline the role these mechanisms may play in endosperm-based hybridization incompatibilities between *M. guttatus* and *M. luteus*.

Differing epigenetic states between parental genomes can lead to deregulation of small RNAs, causing changes in gene body and TE methylation that impact gene expression and phenotype (Rigal et al. 2016; Lafon-Placette and Köhler 2015; Shen et al. 2012). In our system, we find clear differences between the methylation patterns of *M. luteus* and *M. guttatus* endosperm as well as changes in methylation following hybridization. *M. luteus* gene bodies have greater CHG and CHH methylation, while *M. guttatus* has greater CpG methylation on gene bodies and methylation of all types on TEs. Within the hybrids, TE methylation tends to be lower, and CHG and CHH gene body methylation patterns show non-reciprocal patterns depending on their parent of origin. CpG gene body methylation, which is typically either lowly or very highly methylated in the parents (ie. bimodal), loses this pattern in the hybrids. Not only does the amount of TE methylation differ between the parental genomes, but previous work has also shown that the *M. luteus* genome is less TE dense than the *M. guttatus* genome (Edger et al. 2017). Conflicting interactions may especially be prominent between TEs where parents have diverged TE arrangements and methylation states (Senerchia, Felber, and Parisod 2015). Such differences in parental epigenetic landscapes may drive the observed epigenetic reprogramming in the hybrids resulting in subgenome expression dominance and interference with or loss of imprinting patterns. Other global shifts in expression patterns, such as maternal effects, can occur in hybrids (Videvall et al. 2016) and interfere with imprinting as well (Florez-Rueda et al. 2016).

In contrast to the stark *M. luteus* expression dominance over *M. guttatus* homologs in hybrid embryo and endosperm (as well as adult tissues (Edger et al. 2017)), overall expression levels are much more similar between the *M. luteus* ‘A’ and ‘B’ subgenomes in both *M. luteus* and the hybrids. Interestingly, in the *M. luteus* endosperm, the specific parental expression bias of hom*e*ologs (defined in Results) from ‘A’ and ‘B’ subgenomes show little correlation, even though this expression bias, when averaged across all genes, is similar between the two subgenomes. This pattern suggests that, instead of global expression dominance of an entire subgenome, the expression of homeolog pairs is stochastically regulated in *M. luteus*. That said, in the hybrids, the *M. luteus* allele exhibits expression dominance over the *M. guttatus* allele. Both subgenome expression dominance and stochastic regulation of homeolog pairs are mechanisms of coping with polyploidy that have been observed in other allopolyploids (J. Wang et al. 2004; Zhao et al. 2017; Pfeifer et al. 2014; Leach et al. 2014); (Mi-Jeong Yoo et al. 2014). It has been posited that the extent of genetic divergence between parents determines whether global subgenome dominance or more stochastic regulation of homeolog pairs will occur in hybrids; with greater genetic divergence resulting in subgenome dominance (Zhao et al. 2017). However, even if two individuals are genetically very similar, epigenetic differences can induce genomic shock in their offspring (Rigal et al. 2016). In this system, *M. luteus* ‘A’ and ‘B’ subgenomes are no less genetically diverged from each other than they are from *M. guttatus (Edger et al. 2017)*, yet their expression patterns are much more similar in the hybrid. Regulatory patterns may be largely shared by these long coexisting *M. luteus* subgenomes, and may be inherited in the hybrid. Therefore, epigenetic/regulatory differences between parents, beyond purely genetic differences, may substantially drive epigenetic reprogramming and subgenome expression dominance in hybrids. While predicting which subgenome will be dominant is less clear, the decreased density and methylation of TEs on *M. luteus* relative to *M. guttatus* may be related to its expression dominance (Woodhouse et al. 2014; Edger et al. 2017).

As discussed above, the sets of imprinted genes differ between *M. guttatus* and *M. luteus*. Furthermore, for a given set of imprinted genes, their homologs in the other species differ in expression patterns. In the hybrids, imprinted genes have unique patterns of parental expression bias, suggesting more complex mechanisms, such as regulatory interactions between parental genomes, may be at play (Mi-Jeong Yoo et al. 2014). While we cannot explicitly test for epistatic interactions involving these genes with our data, this data does indicate that epigenetic DM-like incompatibilities may be occurring. For example, the lack of overlap in imprinted genes may result in the absence or mismatch of an imprinted gene’s interacting component(s). Methylation and regulatory patterns have clearly diverged to some extent between the two parent species’ endosperm. Interestingly, while subgenome expression dominance likely occurs in hybrid embryo, it is even more pronounced in hybrid endosperm. Thus, large-scale epigenetic reprogramming may interact with the diverged epigenetic characteristics of parental endosperm. While epigenetic reprogramming occurs in each hybrid cross, it differs depending on crossing direction, which may explain the non-reciprocal seed phenotypes. What’s more, the role of subgenome expression dominance with crossing direction likely has important implications for parent-of-origin effects in hybrid endosperm. For example, it could create or increase imbalances in the dosage of maternal and paternal alleles and their interacting components non-reciprocally. The acquired subgenome expression dominance of *M. luteus* PEGs in the hybrid endosperm may indicate such a pattern. However, genetic incompatibilities and more complex mechanisms not assessed here may certainly be involved, and functional studies are needed to test the role different imprinted genes have in these incompatibilities.

Based on the above ideas, we suggest that the following processes may contribute to hybrid barriers in *M. luteus* x *M. guttatus* crosses: (1) Different sets of imprinted genes in parents generate epigenetic DM-like incompatibilities; (2) Disparity of epigenetic characteristics between the parent species drives an epigenomic shock in the hybrids; (3) Methylation changes and other epigenetic reprogramming following hybridization may alter regulatory networks between the parent genomes, whereas epigenetic similarities between the two *M. luteus* subgenomes are maintained; and (4) Global subgenome expression dominance in hybrid endosperm, where the *M. luteus* genome is more expressed, exacerbates imprinting incompatibilities between the two species’ genomes resulting in seed development abnormalities. Given this logic and observations from other systems, we suggest that epigenetic repatterning driving global shifts in expression patterns may result from diverged epigenetic and regulatory landscapes of parental genomes. This may either establish or exacerbate incompatible interactions between specific imprinting patterns in parental species that have diverged in status or dosage (or both) from the ancestral state. Importantly, even if imprinting patterns are mostly conserved, epigenetic differences that shift expression patterns could affect dosage interactions of imprinted genes and produce non-reciprocal incompatibilities. Such a process could serve as a general underlying mechanism for understanding the role of genomic imprinting in endosperm-based hybrid incompatibilities. Since such incompatibilities may likely proceed those in the embryo as well as genotype divergences, this mechanism may be an important component of hybridization barriers in general. Further studies in this system and the many other intriguing *Mimulus* hybrid and speciation systems (Garner et al. 2016; Kooyers, James, and Blackman 2017; Oneal et al. 2014; Hall and Willis 2006; L. Fishman and Willis 2001; Lila Fishman, Kelly, and Willis 2002; Oneal, Willis, and Franks 2016) along with its genomic and genetic resources (Edger et al. 2017; M. Vallejo-Marin et al. 2016; Ding and Yuan 2016) will make *Mimulus* a valuable model for elucidating the mechanisms and evolutionary drivers of genomic imprinting.

## METHODS

### Crossing design

Four accessions were used: CS and Mll for *M. luteus* and CG and LCA for *M. guttatus*. LCA is from Lake Alamanor, California and is 11 generations inbred, and Mll is from Embalse el Yeso, Chile, and is 13 generations inbred. For more information on CS and CG, refer to Vallejo-Marin et al. 2016. For the histology, seed area, and germination experiments, first or second generation CS and CG plants were used. Flowers were emasculated before blooming, and then pollinated by a flower from a different plant the next day. There were four unique crosses in total: *4x x 4x* (*M. luteus*; CS pollinated by CS), *2x x 2x* (*M. guttatus*; CG pollinated by CG), *2x x 4x* (CG pollinated by CS), and *4x x 2x* (CS pollinated by CG). For RNA-Seq, Mll, CS, LCA, and CG were used to produce six unique crosses. There were two reciprocal crosses for *M. luteus*, *Mll x CS* and *CS x M11*, two for *M. guttatus*, *CG x LCA* and *LCA x CG*, and the two hybrid reciprocal crosses, *CG x M11* (*2x x 4x*) and *M11 x CG* (*4x x 2x*). There were four crosses used for Methyl-Seq: *M11 x CS*, *CG x LCA*, *CG x M11*, and *M11 x CG*. Seeds were always collected at the same time of day for consistency. All plants were grown in the College of William and Mary greenhouse with 16 hours of light per day.

### Seed area and histology

Four mature ovaries with 100-200 seeds each were collected for each cross (refer to Crossing design section) at 15-18 days after pollination (DAP). Images were taken of a subset of seeds (used below in the Seed germination section) under a dissecting microscope, with the same magnification for all images, and area and aspect ratio was measured in ImageJ with a custom script to automate measurements. ANOVAs with post-hoc Tukey-Kramer tests were used to compare crosses in R.

To investigate seeds throughout their development, three ovaries from each cross were collected at 3, 5, 8, and 11 DAP (48 ovaries total). Ovaries were immediately placed into 4% paraformaldehyde and vacuum infiltrated for 15-20 minutes and kept at 4C for 48 hours. They were then washed with Phosphate Buffered Saline (PBS) for 30 minutes three times and left in fresh PBS at 4C overnight. Ovaries were then dehydrated using the following dehydration series: 10% ethanol for 15 minutes, 30% for 15 minutes, 50% for 15 minutes, 70% for 15 minutes, 95% for 20 minutes, and two times of 100% for 30 minutes, all on a shaker. Ovaries were next infiltrated through the following infiltration series: 100% propylene oxide three times for 20 minutes each, 2:1 propylene oxide to Spurr’s resin mixture (EMS Catalog #14300) for one hour, and 1:1 propylene oxide to Spurr’s for one hour, all on a shaker. They were embedded into full resin, changed after two hours, and left overnight in fresh resin at room temperature on a shaker. The next day, resin was changed twice every two hours. Ovaries were then placed into individual molds with resin and left in an oven at 60C for 48 hours. Using an ultramicrotome, 0.5 micrometer sections were produced from the resin molds. Sections were adhered to slides in water at 50C on a heat block. Slides were next stained with Azure II for 5 minutes at 50C on a heat block. Coverslips were mounted onto slides using acrylamide, and images were taken on a compound microscope using SPOT Imaging™ software. Sections that contained the center of the seed (for consistency among seeds) were selected from each cross for 8 DAP and 11 DAP. ImageJ was used to measure the area of the endosperm, embryo, and whole seed by manually tracing these tissues. Measurements were adjusted based on the magnification of the image. ANOVAs with post-hoc Tukey-Kramer tests were used in R to compare endosperm and embryo area among crosses at 8 and 11 DAP each. Student’s t-tests were used to identify shifts in area from 8 to 11 DAP for endosperm and embryo.

### SEM

Ten mature seeds (above) from each of the four crosses were fixed in 2% glutaraldehyde with PBS for two hours. After fixation, seeds were washed with PBS and put through a dehydration series at concentrations of 50%, 70%, 85%, 95%, and 100% ethanol:PBS for one hour at each step. Seeds were left in 100% ethanol overnight. One hour before critical point drying, 100% ethanol was renewed. After critical point drying with a Samdri® PVT-3B Critical Point Drier (Tousimis® Research Corporation), seeds were sputter-coated with gold-paladium using a Hummer® sputtering system from Anatech Ltd. and mounted on pin stubs for Scanning Electron Microscopy (SEM). All images were acquired using Phenom Tabletop® Scanning Electron Microscope.

### Seed germination

To link the observed seed morphologies to germination rates, a germination experiment was performed in which morphology was measured. Mature ovaries were collected at 15 - 18 DAP within a week of the experiment (these were the same mature ovaries as in the Seed area and histology section above). Seeds were soaked in 3% calcium hypochlorite for 10 minutes and rinsed in PBS three times for 5 minutes, all on a shaker. They were then placed on 60 mm petri dishes with gridded filter paper atop additional filter paper to retain moisture. The petri dishes and filter paper had been sterilized with UV for 30 minutes first. Images of seeds on the plates were taken with SPOT Imaging™ software, area was measured in ImageJ using a custom script to automate measurements, and seed germination was tracked specifically for each seed based off its position on the grid. 5 plates were used per cross with 16-32 seeds per plate. Seeds were placed under growth lights for 8 days with 16 hours of light each day. The final data sheet contained a list of seeds, their size, which cross they belonged to, whether or not they germinated over the 8 day period, and if so, then on which day they germinated. Student’s t-tests were used in R to compare the area of germinated seeds to non-germinated seeds for seeds from all crosses, seeds from all crosses excluding *4x x 2x*, and seeds from each individual cross. Next, ANOVAs were used in R to compare the area of seeds at different germination dates (2, 3, 4, 5, 6, and 7 days after the start of the experiment) for all crosses and for each individual cross. Finally, since we could not measure endosperm area for this germination experiment, a linear regression was performed in R between endosperm and seed area using the histology data above for 8 and 11 DAP in order to link whole seed area to endosperm area.

### RNA-Seq data collection

Three or four ovaries were collected at 11 DAP for each cross (10 DAP for *M11 x CS*, since little endosperm was left at 11 DAP). After collection, ovaries were stored in RNA*later®* (Ambion®) at 4C. In order to separate endosperm tissue from the rest of the seeds, a modified protocol was used from (Gehring, Bubb, and Henikoff 2009). Endosperm and embryo were dissected from seeds in a 0.3 M sorbitol/5mM MES (2-[N-morpholino] ethanesulfonic acid) solution (Gehring, Bubb, and Henikoff 2009) on a glass slide under a dissecting microscope using 30-gauge hypodermic needles. Endosperm and embryo from 20-40 seeds were pooled (separately) per replicate (an ovary from one pollination event) and rinsed with the same solution 5-10 times. Pooled endosperm and embryo were then placed into the Lysis solution of RNAqueous®-Micro Total RNA Isolation Kit (Ambion®) and disrupted with a pestle (rotating 50-60 times). The remainder of the RNA extraction protocol from the kit was performed. RNA was converted into cDNA and libraries were constructed using KAPA Stranded mRNA-Seq Kit. During library construction, sequence specific Illumina® TruSeq adapters were added to distinguish each library. Using an Agilent 2100 Bioanalyzer, average fragment lengths were determined to be between 250 and 300 bp. Libraries were then pooled and sequenced by the Duke Center for Genomic and Computational Biology on an Illumina® HiSeq 2500 instrument.

### Analysis of RNA-Seq data

Parental genomes were SNP corrected by first mapping whole genome (CG and CS) and transcriptome data (LCA) to previously assembled reference genomes (Edger et al. 2017; Hellsten et al. 2013) using bowtie2 in –very-sensitive-local mode (Langmead and Salzberg 2012). Picardtools were used to fix mate information for paired end reads (‘FixMateInformation’), remove PCR duplicates (‘MarkDuplicates’), and add read groups (‘AddOrReplaceReadGroups’) (https://broadinstitute.github.io/picard). Sequence variants were then called using the GATK UnifiedGenotyper (McKenna, et al. 2010) with parameters: “-R genome -T UnifiedGenotyper -rf MaxInsertSize –maxInsertSize 10000 -rf DuplicateRead -rf BadMate -rf BadCigar –minbasequalityscore 25 -rf MappingQuality –minmappingqualityscore 25 -ploidy 2 –genotypelikelihoodsmodel BOTH –outputmode EMITALLSITES –maxalternatealleles 2 $inputBams -o $outputVCF”. The GATK FastaAlternateReferenceMaker (McKenna et al. 2010) was used to generate the new SNP corrected fasta files. For each cross (refer to the Crossing design section above), the SNP corrected coding regions of the parental genomes were combined into a single fasta file, and the fastq reads were aligned to these references using bowtie2. For aligning RNA-seq reads to the combined reference, the following settings were used: “--local -5 10 -D 25 -R 4 -N 0 -L 10 -i S,1,0.5 --mp 46,42”. Counts were generated using HTSeq-count with the default options (Anders, Pyl, and Huber 2015). Homeologs with no sequence differences were excluded from further analysis. Here, homeologs are the two alleles (ie. maternal and paternal) identified on the two inbred lines’ genomes used in the given cross.

Next, parental allelic expression biases (AEB) and shifts in AEB (AEBS) were calculated and compared between crosses using the methods described in Smith, et al. 2017. Maternal bias has a negative AEB, paternal bias has a positive AEB, and AEB of 0 indicates no bias. For every homeolog pair within each cross, a likelihood ratio test was used to test whether, after normalizing for gene length and sequencing depth differences, the mean expression level of the two homeologs was the same or different, assuming the mean expression level follows a negative binomial distribution. False discovery rates for all tests were controlled using the R package ‘fdrtool’ (Strimmer, 2008), and AEB / AEBS values with a q-value less than 0.05 were called significant. Inspection of unadjusted p-value distributions revealed one case, the embryo of the CG x LCA (*M. guttatus*) cross, with a highly conservative (a theoretically impossible) skew. This might be indicative of unknown covariates in the data, or an otherwise misspecified null model. To correct this, we ran DESeq2 (Love, Huber, and Anders 2014) for each gene (as maternal vs. paternal) and extracted the shrunken log_2_ fold changes as well as their estimated standard errors, then performed a Z-test between each homeolog pair. This method also produced p-values with a highly conservative skew, however fdrtool was able to correct the Z-scores using its empirical null modeling approach. For AEB tests, genes were filtered out if they had less than 10 RPKM in either the paternal or maternal allele, or had at least one replicate with no counts. This test was performed for every gene in *M. guttatus* (ie. CG x LCA and LCA x CG), *M. luteus* (ie. CS x Mll and Mll x CS), and the hybrid (ie. CG x Mll and Mll x CG). For AEBS tests, the same filter was extended to include both reciprocal crosses. Imprinted genes were those that consistently had significant (after controlling FDR<0.05) maternal or paternal bias between the reciprocal crosses (MEG or PEG, respectively). Due to the reciprocal crossing design used (eg. CG x LCA and LCA x CG), genes can either be consistently biased to the maternal or the paternal allele, or consistently biased to either one of the inbred lines (or have no bias in at least one cross). The percentage of this subset of genes (with any bias) that fell into each one of these four categories was calculated. AEB values of genes in one cross were plotted against the AEB values of the same genes in the reciprocal cross (the direction of parental bias is reversed for visualization) for *M. guttatus* (ie. CG x LCA and LCA x CG), *M. luteus* (ie. CS x Mll and Mll x CS), and the hybrid (ie. CG x Mll and Mll x CG). Finally, a Fisher’s test was used to compare the PEG:MEG ratio between *M. luteus* and *M. guttatus*.

The following statistical analyses apply only to endosperm and were performed in R. We first quantified the amount of overlap in imprinted genes between *M. guttatus* and *M. luteus*. The list of genes for the *2x x 4x* and *4x x 2x* crosses represent all the known clear homeologs between *M. guttatus* and *M. luteus* (Edger et al. 2017). Therefore, in order to assess overlap between *M. guttatus* and *M. luteus* imprinted genes, we counted imprinted genes within this list of homologs. Of its 276 putative imprinted genes, *M. luteus* had 38 PEGs with clear homologs in *M. guttatus*, and 1 of *M. guttatus*’ 16 MEGs and 5 of its 37 PEGs had homologs in *M. luteus*. Note that each of these genes in *M. guttatus* has two corresponding homologs in *M. luteus* since it is a tetraploid. Next we wanted to understand how the parental expression bias (ie. AEB) of one species’ imprinted genes differs in the *other* crosses. We focused only on PEGs. There were a total of 46 *M. luteus* and *M. guttatus* homologs that were PEGs in at least one of the species. We gathered AEB data from each cross for this list of genes. For each gene, AEB values were averaged between reciprocal crosses for *M. guttatus* and for *M. luteu*s (eg. *CG x LCA* and *LCA x CG*). We next measured the mean AEB of the 5 *M. guttatus* and 38 *M. luteus* PEGs represented in this list, the mean AEB of those same genes in the *2x x 4x* and *4x x 2x* crosses, the mean AEB of the *M. guttatus* homologs to the *M. luteus* PEGs (there was not enough data for *M. luteus* homologs to the *M. guttatus* PEGs), and the mean AEB of all *M. guttatus* and *M. luteus* homologs represented in each cross. Genes were filtered by low RPKM as stated above for RPKM filtration when considering reciprocal crosses. For *2x x 4x* and *4x x 2x* crosses, genes were filtered out if they had less than 10 RPKM on both the maternal *and* paternal allele and if the number of reads in any replicate was 0. Finally, using the full list of all *M. guttatus* and *M. luteus* homologs, a linear regression was calculated between the two species AEB values. The AEB values used for this regression were the average of the two reciprocal crosses for each species. If in *both* species, a gene’s RPKM was low (as described above when considering reciprocal crosses), it was filtered out.

We next compared the log RPKM of parental alleles in the embryo and in the endosperm for the parental species and the hybrid crosses. For *M. guttatus* and *M. luteus*, the log 10 of the RPKM, averaged between reciprocal crosses, was taken for each allele (maternal and paternal) from each tissue. For *2x x 4x* and *4x x 2x*, the log 10 RPKM was taken for each allele from each tissue (except the embryo of *4x x 2x*). ANOVAs with subsequent pairwise Tukey’s HSD tests were performed comparing the maternal embryo allele, the paternal embryo allele, the maternal endosperm allele, and the paternal endosperm allele for *M. guttatus*, *M. luteus*, and *2x x 4x*. A student’s t-test was used to compare the maternal and paternal alleles in the endosperm for *4x x 2x*. No filtration criteria were used.

### Analysis of Methyl-Seq data

Endosperm was collected and pooled using the same methods as above (see RNA-Seq data collection section for details), and DNA was extracted using a CTAB based protocol. There were 3 replicates for each of the 4 crosses used (refer to the Crossing design section above). Bisulfite conversion was performed using the EZ DNA Methylation™ Kit (Zymo) and libraries were immediately constructed using the TruSeq DNA Methylation Kit (Illumina®) with adapters from TruSeq DNA Methylation Index PCR Primers (Illumina®). DNA was sequenced on an Illumina® HiSeq 2500 generating 50 bp reads. Methylation analyses were conducted using the methylpy pipeline (Schultz et al. 2015). Methylpy was specifically designed to analyze high-throughput bisulfite sequencing data. Methylation for each biological replicate was aligned to a combined reference file using this pipeline with the following parameters: “num_procs=20, illumina adapter sequence, quality_version=“1.8”, bowtie_options=[“-S”,“-k 1”,“-m 1”,“--chunkmbs 3072”,“--best”,“--strata”,“-o 4”,“-e 80”,“-l 20”,“-n 0”], max_adapter_removal=None, overlap_length=None, zero_cap=None, error_rate=None, min_qual_score=10, min_read_len=30, sig_cutoff=0.05, min_cov=3, binom_test=True, num_reads=-1”. The combined reference file contained both maternal and paternal genomes as well as the mitochondrial genome (NCBI accession NC_018041). Following alignment, methylpy calls methylated bases based on SNPs relative to the reference. The output of methylpy includes, for each site, the number of methylated bases sequenced, the total number of bases sequenced, and the type of site (CpG, CHG, CHH). This data in combination with the genome annotation files were used to analyze the data by gene, TE, and methylation type. Nonconversion rates per sample ranged from 0.5% to 4.8%. Counts for each of the three replicates were added together for each cross. After summing, genes were filtered out if the total number of reads mapped to them was 20 or less. For each methylation type, the fraction of methylation on each allele (maternal and paternal) was calculated for each cross and Kolmogorov–Smirnov (KS) tests were performed on the fraction of methylation between all combinations of alleles. The empirical cumulative distribution function was executed and plotted using the ggplot2 package in R for each methylation type. While parental alleles could not be distinguished for TEs, total reads per TE were summed across the three replicates (and filtered as above), the fraction of methylation on each TE was calculated, and KS tests were performed between all combinations of the four crosses for each methylation type. Empirical cumulative distributions were also plotted as above.

### Analysis of subgenome data

Log RPKM was measured in the same way as above, except each *M. luteus* subgenome (A and B) was measured separately in the *M. luteus*, *2x x 4x* and *4x x 2x* crosses (for lists of genes within *M. luteus* subgenomes refer to (Edger et al. 2017)). ANOVAs with subsequent pairwise Tukey’s HSD tests were performed as well, treating *M. luteus* subgenomes for each parental allele and each tissue as different groups.

To compare the relationship of AEB between homeolog pairs (homologous genes between the two subgenomes) in *M. luteus*, a linear regression was performed between AEB values calculated from comparing the maternal and paternal alleles of the A subgenome and the corresponding AEB values calculated from comparing homeologs on maternal and paternal alleles of the B subgenome. In other words, for each homeolog pair, parental expression bias between A subgenome alleles was compared to parental expression bias between B subgenome alleles. Homeolog pairs were filtered out if the number of reads in any replicate was 0 for either homeolog. We did not filter for low RPKM, because sample sizes were low for the linear regressions, though even after filtering out genes that had less than 10 RPKM on both the maternal *and* paternal allele (for each subgenome), results of linear regressions were very similar. Linear regressions were performed for each reciprocal cross (ie. Mll x CS and CS x Mll) in the embryo and the endosperm. Similar methods and filtration were used for the hybrid crosses (ie. *2x x 4x* and *4x x 2x*) except linear regressions were performed between AEB values calculated from comparing the *M. guttatus* (maternal for *2x x 4x* and paternal for *4x x 2x*) and A subgenome alleles (paternal for *2x x 4x* and maternal for *4x x 2x*) and the corresponding AEB values calculated from comparing the *M. guttatus* and B subgenome alleles. Linear regressions were performed in the embryo and endosperm of *2x x 4x* and the endosperm of *4x x 2x*. Again genes were not filtered for low RPKM, but even if filtered, results were also similar.

### BLAST of imprinted genes

Using a list of differentially imprinted genes we took the gene name and found the sequence from a specific reference genome (either *M. luteus* or *M. guttatus).* This reference sequence was then BLASTed against a database of *Arabidopsis thaliana* genes (TAIR). We used a BLAST NT query and set outputs to include Araport 11 Transcripts (DNA). Gene descriptions were pulled from *A. thaliana* genes with e-values below 0.01.

## ACKNOWLEDGEMENTS

This work was supported by The College of William and Mary Research Award to J.R.P. We would like to thank Dr. Diane Shakes and Dr. Harmony Dalgleish for their support and constructive critique during the development and implementation of this project. Dr. Abigail Reft provided support and training for seed sectioning and SEM. Colleen Flynn performed SEM. Hunter Call wrote scripts for automated measurements of seed morphology in ImageJ. Sara Meier, Nora Flynn, and Scott Teresi assisted in lab work. We would also like to thank Dr. Mary Gehring for providing insight into seed dissection methods for RNA extraction.

## AUTHOR CONTRIBUTIONS

T.J.K. and J.R.P. conceived of and designed all aspects of this research. A.M.C and M.V. contributed initial ideas for research development, edited this article and its conclusions, and provided seeds. T.J.K. performed all crosses for histology, germination experiments, and sequencing; performed seed sectioning, staining, histological measurements, germination experiments, and all associated data analyses; collected samples and performed RNA sequencing and bisulfite treatment/Methylation sequencing. R.D.S. performed transcriptome assembly and all homeolog expression analyses and likelihood ratio tests with input from J.R.P and T.J.K. R.D.S. and G.D.S. modified these analyses for this system and the specific questions addressed in this study. Furthermore, R.D.S. provided general guidance regarding statistical analyses. A.H.L. performed methylation analyses with input from J.R.P and T.J.K. T.J.K. performed downstream analyses of homeolog expression and methylation data with input from J.R.P. T.J.K. wrote this article with contributions from R.D.S, J.R.P, and A.H.L.

